# Identifying good practices for detecting inter-regional linear functional connectivity from EEG

**DOI:** 10.1101/2022.10.05.510753

**Authors:** Franziska Pellegrini, Arnaud Delorme, Vadim Nikulin, Stefan Haufe

## Abstract

Aggregating voxel-level statistical dependencies between multivariate time series is an important intermediate step when characterising functional connectivity (FC) between larger brain regions. However, there are numerous ways in which voxel-level data can be aggregated into inter-regional FC, and the advantages of each of these approaches are currently unclear.

In this study we generate ground-truth data and compare the performances of various pipelines that estimate directed and undirected linear phase-to-phase FC between regions. We test the ability of several existing and novel FC analysis pipelines to identify the true regions within which connectivity was simulated. We test various inverse modelling algorithms, strategies to aggregate time series within regions, and connectivity metrics. Furthermore, we investigate the influence of the number of interactions, the signal-to-noise ratio, the noise mix, the interaction time delay, and the number of active sources per region on the ability of detecting phase-to-phase FC.

Throughout all simulated scenarios, lowest performance is obtained with pipelines involving the absolute value of coherency. Further, the combination of dynamic imaging of coherent sources (DICS) beamforming with directed FC metrics that aggregate information across multiple frequencies leads to unsatisfactory results. Pipeline that show promising results with our simulated pseudo-EEG data involve the following steps: (1) Source projection using the linearly-constrained minimum variance (LCMV) beamformer. (2) Principal component analysis (PCA) using the same fixed number of components within every region. (3) Calculation of the multivariate interaction measure (MIM) for every region pair to assess undirected phase-to-phase FC, or calculation of time-reversed Granger Causality (TRGC) to assess directed phase-to-phase FC. We formulate recommendations based on these results that may increase the validity of future experimental connectivity studies.

We further introduce the free ROIconnect plugin for the EEGLAB toolbox that includes the recommended methods and pipelines that are presented here. We show an exemplary application of the best performing pipeline to the analysis EEG data recorded during motor imagery.

## 1. Introduction

In recent years, the field of functional neuroimaging has seen a shift from the mere localization of brain activity towards assessing interaction patterns between functionally segregated and specialized brain regions (Friston, 2011; Schoffelen and Gross, 2019). Functional connectivity (FC), in contrast to structural connectivity, expresses a statistical dependency between two or more neuronal time series. It has been proposed that FC reflects inter-areal brain communication (Fries, 2015). Moreover, empirical FC estimates have been linked to various cognitive functions (Schoffelen and Gross, 2019) and show pathological alterations in many neurological diseases like Parkinson’s Disease, Alzheimer’s Disease, and epilepsy (Van Diessen et al., 2015).

Electroencephalography (EEG) and Magnetoencephalography (MEG) are suitable tools for recording neural activity non-invasively with high temporal resolution. Pipelines for analysing inter-regional FC from M/EEG recordings typically consist of a series of processing steps: artifact cleaning, source projection, aggregation of signals within regions of interests (ROIs), and, finally, FC estimation. At each step, researchers can choose between a huge selection of processing methods, where every decision has the potential to crucially affect the final result of an analysis and its interpretation (Wang et al., 2014; Colclough et al., 2016; Mahjoory et al., 2017). This not only complicates the comparison of results from different FC studies, it also raises the question: which pipelines are suitable for reliable source-level FC detection from M/EEG?

In the absence of a robust ground truth on information flow patterns in the human brain, computer simulations are a straightforward way to address such questions (Ewald et al., 2012). Indeed, numerous works have aimed to validate parts or aspects of M/EEG FC methodologies by employing simulated activity. Several studies have focused on assessing the accuracy of different inverse solutions (Grova et al., 2006; Haufe et al., 2008, 2011; Castaño-Candamil et al., 2015; Bradley et al., 2016; Hincapié et al., 2017; Anzolin et al., 2019; Halder et al., 2019; Jaiswal et al., 2020; Hashemi et al., 2021; Allouch et al., 2022). Others have tested the performance of different FC metrics (Astolfi et al., 2007; Silfverhuth et al., 2012; Haufe et al., 2013; Anzolin et al., 2019; Sommariva et al., 2019; Allouch et al., 2022); however, not always on source-reconstructed data exhibiting realistic levels of source leakage.

Many studies aim at aggregating FC within physiologically defined ROIs (Supp et al., 2007; Palva et al., 2010, 2011; Schoffelen et al., 2017; Basti et al., 2020; Idaji et al., 2021). This approach has various advantages. First, it is computationally more tractable (both memory- and time-wise) than the computation of FC between many pairs of individual sources, and it can avoid numerical instabilities for FC metrics that require full-rank signals. Second, interpreting or even visualizing FC between thousands of separate sources is almost impossible. Third, statistical testing is far easier due to a much reduced number of multiple comparisons. And, forth, across-subject statistical analyses are eased by working on a standardized set of regions rather than in individual anatomical spaces lacking a common set of source locations.

There have been various suggestions on how to reduce the signal dimensionality within ROIs. While some approaches focus on selecting one source for each ROI that best represents the activity of all sources in it (Hillebrand et al., 2012; Ghumare et al., 2018; Perinelli et al., 2022), others involve some kind of averaging or weighted averaging over all source time series of a ROI (Palva et al., 2010, 2011; Korhonen et al., 2014). This approach can be made more general by using the strongest principal component (PC) of all sources of a ROI as a representative time series of that ROI (Supp et al., 2007; Hillebrand et al., 2012; Ghumare et al., 2018; Rubega et al., 2019; Basti et al., 2020). The assumption behind this is that the projection of the data that captures the highest amount of variance within a ROI (its strongest PCs) also reflects the connectivity structure of that ROI best. While most works use only the first PC per region, the use of multiple components has also been suggested (e.g. Schoffelen et al., 2017). For this approach, the subsequent FC estimation is usually calculated between pairs of multivariate time series. Another approach, used for example in Schoffelen et al. (2017), is to apply a multivariate FC metric (here, a multivariate extension of Granger causality, Barrett et al., 2010) to the first *C* PCs of each pair of ROIs. Comparable undirected metrics are the multivariate interaction measure (MIM) and the maximized imaginary coherency (MIC) (Ewald et al., 2012; Basti et al., 2020), which are currently already in use for source-to-source FC estimation (e.g. D’Andrea et al., 2019). These are promising approaches towards more reliable FC estimation. But their virtue in the context of inter-regional FC estimation is still unclear. Moreover, a comprehensive approach evaluating entire data analysis pipelines rather than individual steps is still lacking (see Mahjoory et al., 2017; Haufe and Ewald, 2019).

Consequently, this work addresses the following questions: First, which pipelines are promising candidates for inferring phase-to-phase FC? Second, which pipelines are promising candidates for inferring the directionality of an interaction? And, most importantly, which pipelines are not suitable to detect FC from data that is corrupted by signal mixing? In addition, we investigate how the number of PCs per ROI affects FC estimation. Finally, we evaluate how the performance of detecting ground-truth interactions varies depending on crucial data parameters like the signal-to-noise ratio (SNR), the number of ground-truth interactions, the noise composition, and the length of the interaction delay. All pipelines are tested within an EEG signal simulation framework that builds on our prior work (Haufe and Ewald, 2019). Note that we focus here on 1:1-phase-to-phase coupling with non-zero time delay, which is the most commonly studied type of FC. Other coupling types including phase–amplitude, amplitude– amplitude, phase–frequency, frequency–frequency, and amplitude–frequency coupling (e.g., Jirsa and Müller, 2013) are not studied here.Further note that we do not intend to propose a realistic model of EEG data or the whole brain. Rather, we aim to identify metrics and pipelines that can accurately reconstruct ROI-level functional connectivity (FC) in the presence of signal mixing, which heavily affects popular metrics used to infer directed and undirected linear FC. That is, we don’t address the question of whether networks estimated using FC metrics provide an accurate depiction of actual brain networks.

The best-performing methods and pipelines identified in this study are implemented in the free ROIconnect plugin for the EEGLAB toolbox. We describe the functionality of ROIconnect and apply it to investigate EEG phase-to-phase FC during left and right hand motor imagery.

## 2. Methods

### 2.1. Data generation

We generate time series at a sampling rate of 100 Hz with a recording length of three minutes (*N*_*t*_ = 100 · 60 · 3 = 18000 samples). For spectral analyses, we epoch the data into *N*_*e*_ = 90 segments of *T* = 200 samples (2 seconds) length.

Ground-truth activity of interacting sources (c.f. Figure 1a) is generated as random white noise filtered in the alpha band (8 to 12 Hz). Throughout, we use zero-phase forward and reverse second-order digital band-pass Butterworth filters. The interaction between two regions is modeled as unidirectional from the sending region to the receiving region. This is ensured by defining the activity at the receiving region to be an exact copy of the activity at the sending region with a certain time delay (see Section 3). Additionally, pink (1*/*f scaled) background noise is added to the sending and receiving regions independently. More specifically, both the ground-truth signal and the pink background noise are first normalized to have unit-norm in the interacting frequency band. To this end, every interacting ground truth signal time series 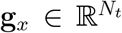 at region *x* is divided by its -norm: 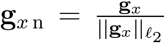.

**Figure 1:**
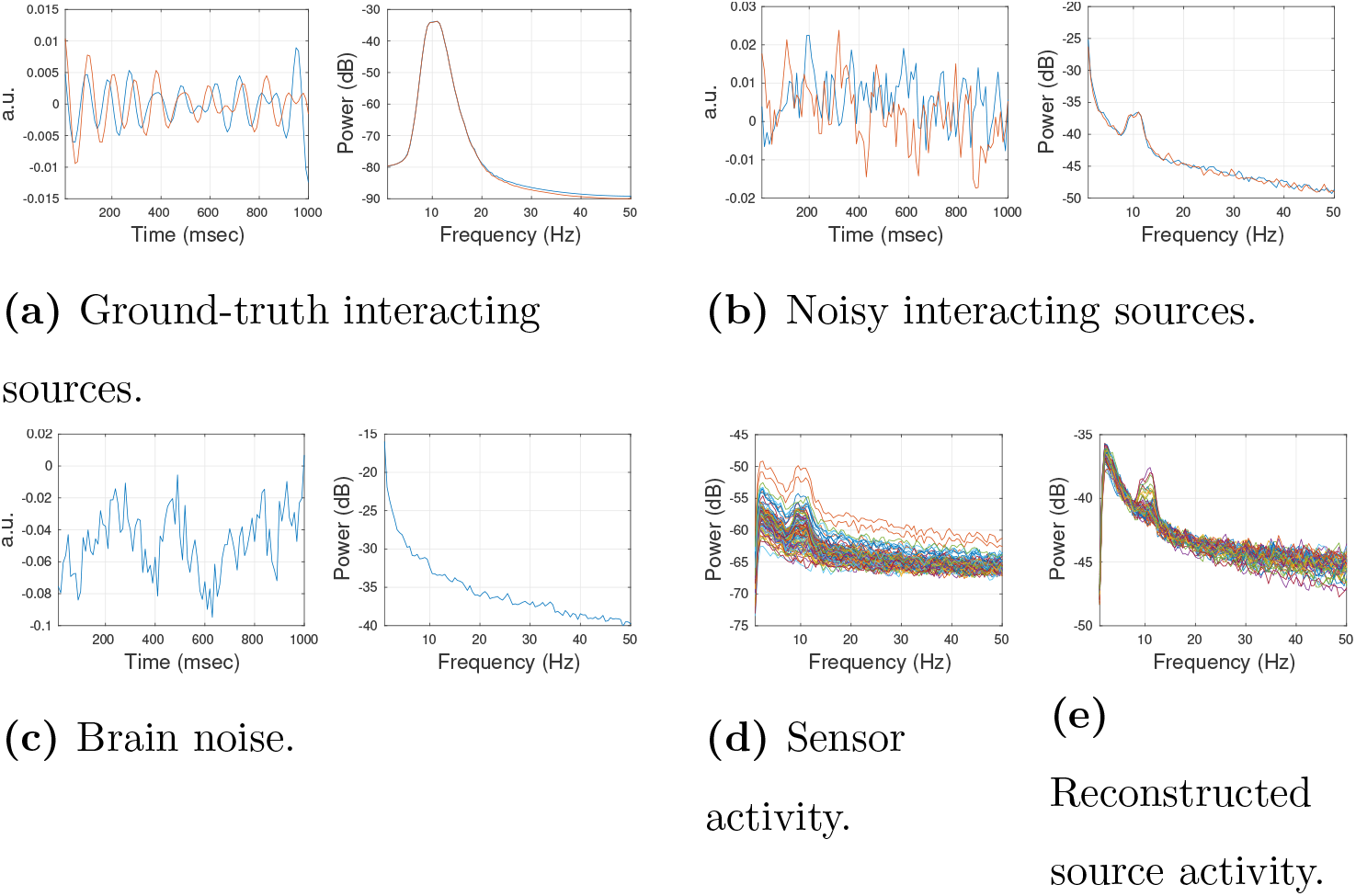
Example of simulated data in time and frequency domain. (a) Ground-truth activity at two interacting sources was generated as random white noise filtered in the alpha band (8 to 12 Hz). Left: the one-second window of data in the time domain. Right: Power spectral density (PSD). (b) Two interacting signals, generated as a mixture of the ground truth activity and pink background noise (SNR_*θ*_ = 3.5 dB). Left: one-second window of data in the time domain. Right: PSD. (c) Brain noise, generated as random pink noise without additional activity in the alpha band (shown is the activity of an exemplary non-interacting source). Left: one-second window of data in time domain. Right: PSD. (d) PSD of activity at the sensor level is generated by mixing white sensor noise, and the interacting signal, and the brain noise at the sensor level (SNR = 3.5 dB). (e) PSD of reconstructed source-level activity. Shown are PSDs of the first principal component of all 68 regions.

Every pink background noise time serie 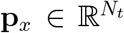 is filtered in the interacting frequency band to obtain 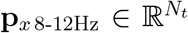. The unfiltered noise time series is then divided by the _2_-norm of its filtered version: 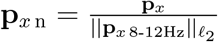.

Subsequently, a weighted sum of the normalized signal time series and the normalized noise time series is calculated:

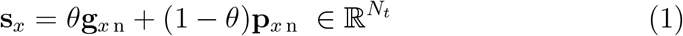

The result is called the (interacting) *signal* (Figure 1b). The parameter *θ* takes values between 0 and 1 and defines the source-level SNR in decibel 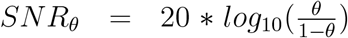. The source-level SNR is set to 3.5 dB (*θ*=0.6). The transposed column vectors of all 2*N*_*I*_ signal time series **s** form the signal sources 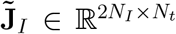, with *N*_*I*_ region pairs containing the 2*N*_*I*_ interacting signals.

In contrast, activity of a non-interacting source at region 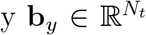 referred to as *brain noise* (Figure 1c) – is generated using random pink noise only without additional activity in the alpha band. The transposed column vectors of all *R −* 2*N*_*I*_ brain noise time series **b** form the brain noise sources 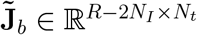, with *R* denoting the number of regions.

We use a surface-based source model with *N*_*v*_ = 1895 dipolar sources placed in the cortical gray matter. Regions are defined according to the Desikan-Killiany atlas (Desikan et al., 2006), which is a surface-based atlas with *R* = 68 cortical regions. Depending on the number of interacting voxels (see Experiment 6, Section 3), one or two time series per region are generated. Every ground-truth time series is placed in a randomly selected source location within a region, so that every region contains the same number of ground-truth time series. The *N*_*I*_ region pairs containing the 2*N*_*I*_ interacting signals are chosen randomly, and all other regions contain time series with brain noise.

In the next step, source activity is projected to sensor space by using a physical forward model of the electrical current flow in the head, summarized by a leadfield matrix. The leadfield describes the signal measured at the sensors for a given source current density. It is a function of the head geometry and the electrical conductivities of different tissues in the head. The template leadfield is obtained from a BEM head model of the ICBM152 anatomical head template, which is a non-linear average of the magnetic resonance (MR) images of 152 healthy subjects (Mazziotta et al., 1995). We use Brainstorm (Tadel et al., 2011) and openMEEG (Gramfort et al., 2010) software to generate the headmodel and leadfield. *N*_*s*_ = 97 sensors are placed on the scalp following the standard BrainProducts ActiCap97 channel setup. Note that the spatial orientation of all simulated dipolar sources is chosen to be perpendicular to the cortex surface, so the three spatial orientations that define the dipole orientation of the source activity orientations are summarized into one. This assumption implies a scalar leadfield 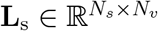. We denote thecolumns of **L**_s_ that correspond to the interacting sources by 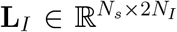 and those corresponding to the brain noise sources by 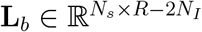. Signal sources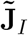 and brain noise sources 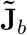 are then separately projected to sensor space:

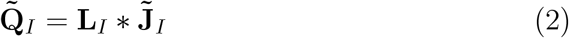

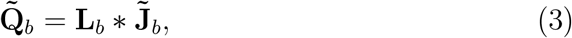

With 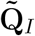 and 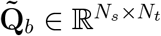.

At sensor level, we mix the different signal and noise components. We generate white sensor noise 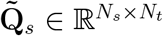 with equal variance at all sensors. The multivariate sensor-space time series corresponding to all three signal components—brain noise, interacting signals, and sensor noise—are divided by their Frobenius norms with respect to the interacting frequency band (8-12 Hz):

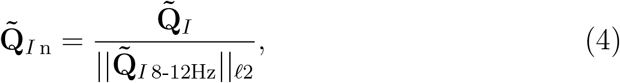

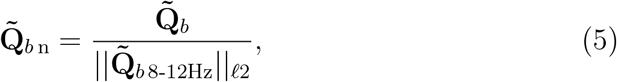

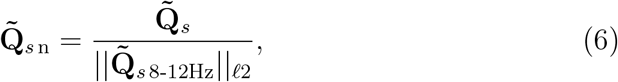

with 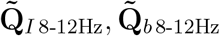 and 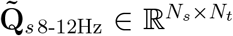. Then the three components are combined as follows: first, we add brain noise and sensor noise with a specific brain noise-to-sensor noise-ratio (BSR) to obtain the total noise 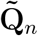 and normalize it with respect to the interacting frequency band:

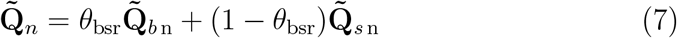

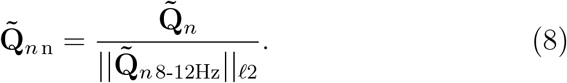

The default BSR value is set to 0 dB, i.e., *θ*_bsr_ = 0.5. Second, we sum up signal and total noise with a specific global (sensor-level) SNR:

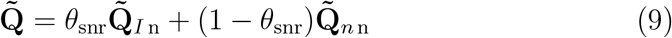

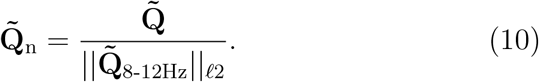

The default SNR value is set to 3.5 dB, i.e., *θ*_snr_ = 0.6. An example of the power-spectral density of the resulting activity on sensor level is illustrated in Figure 1d. As a last step, we high-pass filter the generated sensor data with a cutoff of 1 Hz.

### 2.2. Source reconstruction

We test four different inverse solutions for source reconstruction: ‘Exact’ low-resolution electromagnetic tomography (eLORETA), linearly-constrained minimum variance beamforming (LCMV), dynamic imaging of coherent sources (DICS), and Champagne. Inverse source reconstructions are based on the same leadfield used to simulate the signals. Full 3D currents are estimated for each source dipole. That is, prior information about the dipoles’ orientation is not used. A normal direction could in principle be estimated from the reconstructed cortical surface mesh (which we used here for signal generation); however, such estimation is considered to be rather unstable, since we do not have a good estimate of the cortical surface orientation in practice. The aggregation of the three spatial dimensions is discussed in Section 2.3.

#### ‘Exact’ low-resolution electromagnetic tomography

The starting point to solve the source localization problem is the linear forward model 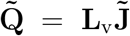, where 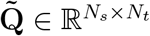 stands for the sensor measurements, 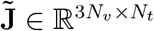 is the vector-valued activity of the dipolar brain sources to be recovered, and 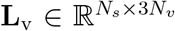 is the vector-valued linear leadfield matrix that maps the electrical activity from sources to sensor level. Here, 3*N*_*v*_ stand for the three spatial dimensions that together define the dipole orientation of the source activity. The solution of this equation is ill-posed since the number of brain sources *N*_*v*_ is much smaller than the number of measurement sensors *N*_*s*_. Therefore eLORETA imposes the constraint of spatially smooth current density distributions (Pascual-Marqui, 2007; Pascual-Marqui et al., 2011). Briefly, eLORETA uses a weighted minimum norm criterion to estimate the source distribution:

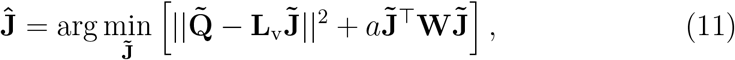

where *a* ≥ 0 denotes a regularization parameter, and **W** is a block-diagonal symmetric weight matrix:

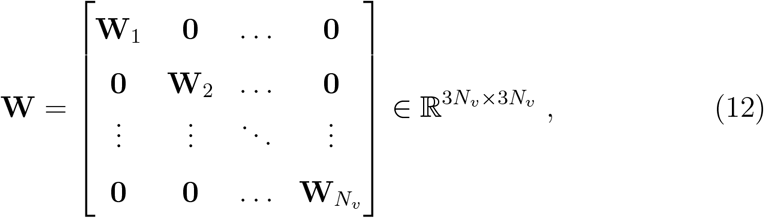

where **0** is the 3 × 3 zero matrix and **W**_*v*_ the 3 × 3 weight matrix at the *v*-th voxel defined in Equation (15). The solution of Equation (11) is given by:

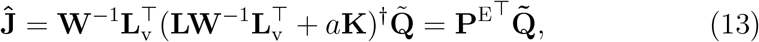

where 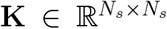 is a centering matrix re-referencing the leadfield and sensor measurements to the common-average reference, *A*^*†*^ is the Moore-Penrose pseudo-inverse of a matrix *A*, and 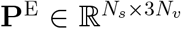 is the eLORETA inverse filter. eLORETA then first computes

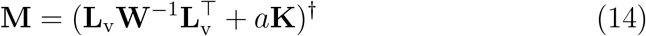

and then for *v* = 1, …, *N*_*v*_, calculates weights

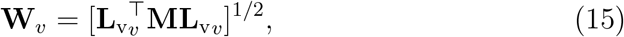

with 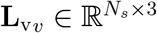 denoting the leadfield for a single source location. It then iterates Equation (14) and (15) until convergence and use the final weights to calculate 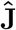. eLORETA has been shown to outperform other linear solution in localization precision (Pascual-Marqui, 2007; Halder et al., 2019; Allouch et al., 2022).

In this study, we choose the regularization parameter based on the best result in a five-fold spatial cross-validation (Hashemi et al., 2021) with fifteen candidate parameters taken from a logarithmically spaced range between 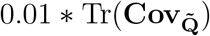 and 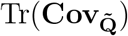, where Tr(*A*) denotes the trace of a matrix *A* and 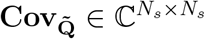 denotes the sample covariance matrix of the sensor-space data.

#### Linearly-constrained minimum variance beamforming

The LCMV (Van Veen et al., 1997) filter 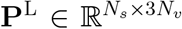 belongs to the class of beamformers. It estimates source activity separately for every source location. While LCMV maximizes source activity originating from the target location, it suppresses noise and other source contributions. Let 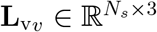 and 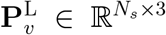 denote the leadfield and projection matrix for a single source location, respectively. The LCMV projection filter minimizes the total variance of the source-projected signal across the three dipole dimensions:

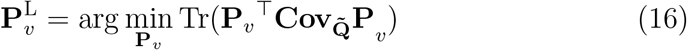

under the unit-gain constraint

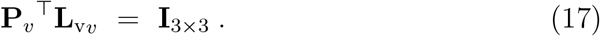

The source estimate 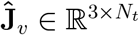 at the *v*-th voxel is given by

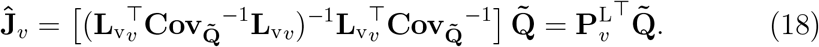

Previous simulations indicated that LCMV overall shows a higher connectivity reconstruction accuracy than eLORETA but is more strongly affected by low SNR (Anzolin et al., 2019). We show a power spectrum of exemplary LCMV-reconstructed source activity in Figure 1e.

#### Dynamic imaging of coherent sources

DICS (Gross et al., 2001) is the frequency-domain equivalent of LCMV. In contrast to LCMV, DICS estimates spatial filters separately for each spectral frequency. The DICS filter **P**^D^ is evaluated for a given frequency *f* using the real part of the sensor-level cross-spectral density matrix **S**_**Q**_:

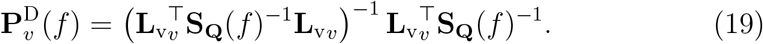

with

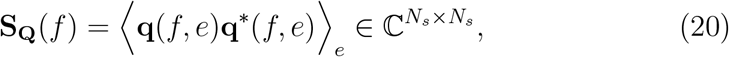

where (·)^*\**^ denotes complex conjugation and **q**(*f, e*) denotes the Fourier transform of the sensor measurements 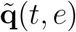.That is, the time-domain sensor signal 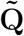 is cut into *N*_*c*_ epochs of *T* time samples to derive 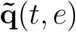, then multiplied with a Hanning window of length *T*, and Fourier-transformed epoch by epoch to derive **q**(*f, e*).

The beamformer filter 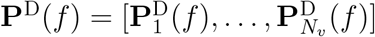 can then be used to project the sensor cross-spectrum to source space:

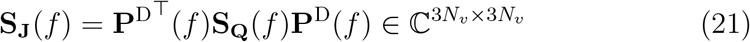

Based on previous literature described above, we hypothesize that the beamformer solutions (LCMV and DICS) perform better than eLORETA when used in combination with undirected FC measures. However, since directed FC measures need to aggregate information across frequencies, we hypothesize that the estimation of such measures might be negatively affected by DICS source reconstruction. Concretely, we expect that DICS’ ability to optimize SNR per frequency and, thereby, to reconstruct different sources for each frequency can be counterproductive in cases where in fact the same pairs of sources are interacting at multiple frequencies. In contrast, we expect that LCMV, which reconstructs a single set of sources by optimizing the SNR across the whole frequency spectrum, would yield more consistent source cross-spectra and, therefore, better directed FC estimates than DICS.

#### Champagne

Champagne (Wipf et al., 2010) uses hierarchical sparse Bayesian inference for inverse modelling. Specifically, it imposes a zero-mean Gaussian prior independently for each source voxel. The prior source covariance is given by

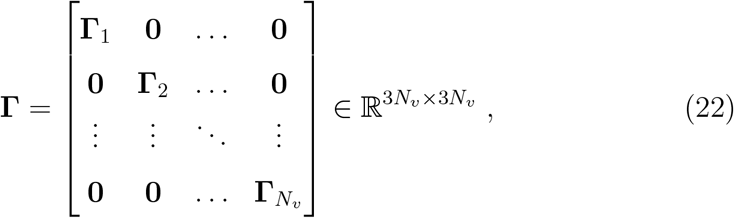

where **Γ**_*v*_ is the 3 × 3 covariance of the *v*-th voxel. Here we use a Champagne variant that models each **Γ**_*v*_ as a full positive-definite matrix

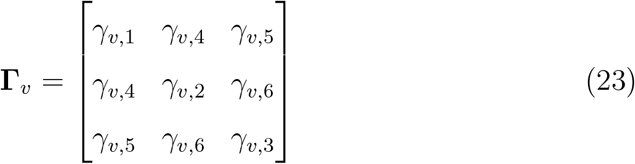

with six parameters. The prior source variances and covariances in **Γ** are treated as model hyperparameters and are optimized in an iterative way.

For any given choice of **Γ**, the posterior distribution of the source activity is given by (Wipf et al., 2010):

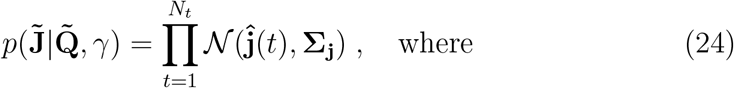

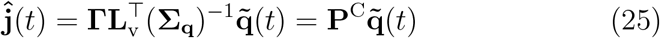

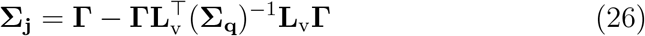

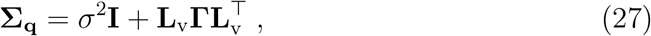

and where *σ*^2^ denotes a homoscedastic sensor noise variance parameter. The posterior parameters 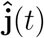 and **Σ**_**j**_ are then used to obtain the next estimate of *γ* by minimizing the negative log model evidence (Bayesian Type-II likelihood):

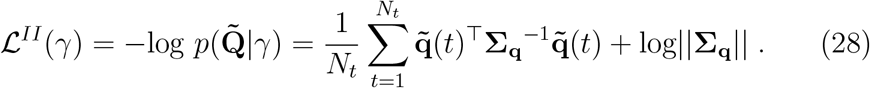

This process is repeated until convergence. Importantly, the majority of source variance parameters converges to zero in the course of the optimization, so that the reconstructed source distribution becomes sparse.

In the original Champagne version, a baseline or control measurement is used to estimate noise covariance in sensor data. Since baseline data are not available in our study, we use a homoscedastic noise model in which all sensors are assumed to be perturbed by uncorrelated Gaussian white noise with equal variance, and estimate the shared variance parameter using five-fold spatial cross-validation (Hashemi et al., 2021). Again, fifteen candidate parameters are taken from a logarithmically spaced range between 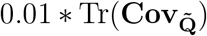 and 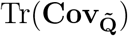.

### 2.3. Dimensionality reduction

To aggregate time series of multiple sources within a region, an intuitive approach would be to take the mean across sources within each spatial dimension. However, this approach has two disadvantages: First, it assumes a high homogeneity within all voxels of a pre-defined region, which is not always given. Second, it does not offer a solution for aggregating the three spatial dimensions, since averaging across these might lead to cancellations due to different polarities.

#### Principal component analysis

An alternative approach is to reduce the dimensionality of multiple time series by employing a singular value decomposition (SVD) or, equivalently, principal component analysis (PCA), and to subsequently only select the *C* strongest PCs accounting for most of the variance within a region for further processing. Let 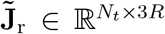 denote the reconstructed broad-band source time courses of *R* dipolar sources within a single region r after mean subtraction. The covariance matrix 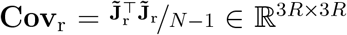 is a symmetric matrix that can be diagonalized as

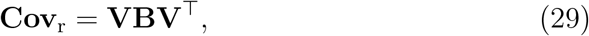

where **B** ∈ ℝ^3*R*×3*R*^ is a diagonal matrix containing the eigenvalues *λ*_*v*_ (variances) of the PCs, which are, without loss of generality, assumed to be given in descending order, and **V** ∈ ℝ^3*R*×3*R*^ is a matrix of corresponding eigenvectors in which each column contains one eigenvector. The *j*^*th*^ PC can then be found in the *j*^*th*^ column of 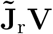.

In practice, the PCs are calculated using an SVD of the zero-mean data matrix 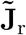 as

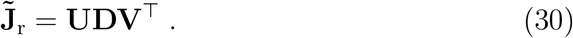

Using the ‘economy version’ of the SVD, 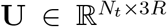 is a matrix of orthonormal PC time courses, **D** *∈* R3*R*×3*R* is a matrix of corresponding singular values, and **V** ∈ ℝ^3*R*×3*R*^ is the matrix of eigenvectors (or, equivalently, singular vectors) defined above. Note that the square of the elements of **D**, divided by *N*_*t*_ − 1 are identical to the variances of the corresponding PCs (eigenvalues of **Cov**_r_). Each squared singular vector, normalized by the sum of all singular vectors, thus corresponds to the variance explained by the corresponding singular vector. We will use this property for the two VARPC pipelines (Section 2.5).

Comparing PCA and SVD, one can easily see that

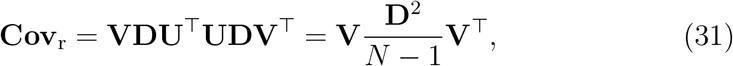

and 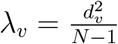.Thus, the PCs can also be calculated with SVD:

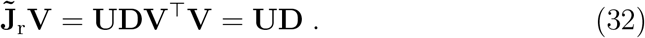

To reduce the dimensionality of the voxel data within one region, we keep only the strongest *C* PCs, i.e., the columns of **UD** that correspond to the largest eigenvalues. For a more extensive overview of the relationship between SVD and PCA, we refer to Wall et al. (2003). Note that in this study, we applied SVD on the time-domain source signals 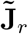 for most of the pipelines. However, we applied PCA on the real part of the source-level cross-spectrum, summed across frequencies, in case of DICS. For the ease of reading, we will stick to PCA terminology for all pipelines in the following.

It has been popular in the literature (Friston et al., 2006; Basti et al., 2020) to select only the first PC for every region and subsequently employ a univariate FC measure for further processing. We describe this approach further in Section 2.5, pipeline FIXPC1.

### 2.4. Connectivity metrics

There are numerous approaches to estimate FC (Schoffelen and Gross, 2019). One key distinction can be made between FC metrics that measure undirected (symmetric) interactions between signals and those that also measure the direction of FC.

It has been shown that the estimation of both undirected and directed FC from M/EEG recordings is complicated by the presence of mixed noise and signal sources (Nolte et al., 2004; Haufe et al., 2013; Bastos and Schoffelen, 2016; Wang et al., 2018; Schaworonkow and Nikulin, 2021). Due to volume conduction in the brain, signal sources from all parts of the brain superimpose at each M/EEG sensor. Projecting the sensor signals to source space can help disentangling separate signal sources. However, a signal reconstructed at a specific source voxel may still contain contributions from other sources in its vicinity. This phenomenon is called source leakage (Schoffelen and Gross, 2009).

Volume conduction and source leakage can lead to spurious FC despite the absence of genuine interactions (Nolte et al., 2004; Haufe et al., 2013).

To overcome this problem, *robust* FC metrics have been developed (Nolte et al., 2004, 2008; Haufe et al., 2013; Winkler et al., 2016). Robustness is here referred to as the property of an FC measure to converge to zero in the limit of infinite data when the observed data are just instantaneous mixtures of independent sources (Nolte et al., 2004). Robust FC metrics use that spurious interactions due to signal mixing are instantaneous, while physiological interactions impose a small time delay. Robust FC metrics are therefore only sensitive to statistical dependencies with a non-zero time delay while eliminating zero-delay contributions.

We here test six different FC measures, four to detect undirected FC (coherence, iCOH, MIC, and MIM), and two measures that estimate the direction of interaction between two sources (multivariate GC and TRGC). This selection includes four robust FC metrics (c.f. Section 1) and two nonrobust ones (coherence and GC). Based on the literature described above, we hypothesize that robust metrics will perform better than non-robust metrics. Please note that all tested FC metrics are frequency-resolved. That is, all metrics output an *N*_*roi*_ × *N*_*roi*_ × *N*_*freq*_ tensor that contains the estimated FC for all region pairs at all frequencies. However, since we expect the interaction to be located in the interacting frequency band between 8 and 12 Hz (see Section 2.1), we select only those frequency bins within this band and average the FC scores across them. As a result, we obtain an *N*_*roi*_ × *N*_*roi*_ matrix.

All tested FC metrics are derived from the cross-spectrum 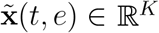and 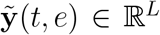 be two multivariate time series where *t ∈* {1, …, *T* } indexes samples within epochs of 2 seconds length and *e* indexes epochs. Often, *K* = *L* = 3 represents the three dipole orientations of two reconstructed current sources. In other cases, *K* and *L* denotes the number of retained data dimensions of two brain regions after (e.g., PCA) dimensionality reduction. These time-domain data are then multiplied with a Hanning window and Fourier transformed into **x**(*f, e*) and **y**(*f, e*), where *f* ∈ {0, 0.5, …, 50} indexes frequencies. The joint cross-spectrum is then computed from the Fourier transformed data as

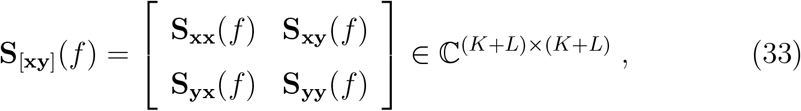

#### Coherence and imaginary part of coherency

(Absolute) coherence (COH) and iCOH are measures of the synchronicity of two time series. Both coherence and iCOH are derived from the complex-valued coherency, which is a generalization of correlation in the frequency domain. As such, coherency quantifies the linear relationship between two time series at a specific frequency. Its phase expresses the average phase difference between the two time series, whereas its absolute value expresses the stability of the phase difference.

Complex-valued coherency **C**_**xy**_ ∈ ℂ^*K*×*L*^ is the normalized cross spectrum (Nunez et al., 1997):

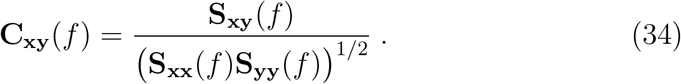

Based on the terminology of Nolte et al. (2004), we define *coherence* as the absolute part of coherency: **COH**_**xy**_(*f*) = |**C**_**xy**_(*f*) | ∈ ℝ^*K*×*L*^, where | · | denotes the absolute value. Coherence captures both zero-delay and non-zero-delay synchronization between two time series. This can be problematic in the context of M/EEG measurements, where substantial zero-delay synchronization can be introduced by signal spread due to volume conduction or source leakage in absence of genuine interactions between distinct brain areas (Nolte et al., 2004). In contrast, the imaginary part of coherency is a robust FC measure since it is only non-zero for interactions with a phase delay different from multiples of *π* (Nolte et al., 2004). Here, we use the absolute value of the imaginary part of coherency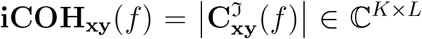, as a measure of synchronization strength, where **C**^ℑ^ denotes the imaginary part of **C**.

Note that both coherence and iCOH are not designed to aggregate FC between two multivariate time series into one FC score. A single FC score can be obtained by taking the average across all elements of **COH**_**xy**_ or **iCOH**_**xy**_, respectively.

#### Multivariate interaction measure and maximized imaginary coherency

The multivariate interaction measure (MIM) and maximized imaginary coherency (MIC, Ewald et al., 2012) are multivariate generalizations of iCOH and are therefore also robust against source leakage.

MIM is defined as follows:

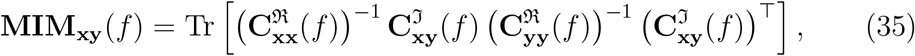

where **C**^**ℜ**^ denotes the real part of **C**. In contrast, MIC aims at maximizing iCOH between the two multivariate time series. That is, MIC finds projections from two multi-dimensional spaces to two one-dimensional spaces such that iCOH between the projected signals becomes maximal:

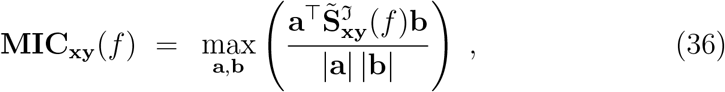

where 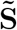 is a whitened version of the cross-spectrum **S** (Ewald et al., 2012), and where **a** *∈* ℝ^*K*×1^ and **b** ∈ ℝ^*L*×1^ are projection weight vectors corresponding to the subspaces, or regions, of **x** and **y**, respectively. Note that, while the imaginary part itself can be positive or negative, flipping the sign of either **a** or **b** will also flip the sign of the imaginary part. Thus, without loss of generality, maximization of Eq. (36) will find the imaginary part with strongest magnitude.

All undirected FC metrics (COH, iCOH, MIC, and MIM) are bounded between 0 and 1.

#### Multivariate Granger causality and time-reversed Granger causality

Granger Causality (GC) defines directed interactions between time series using a predictability argument (Granger, 1969; Bressler and Seth, 2011). Considering two univariate time series 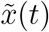 and 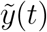, we say that 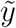 Grangercauses 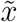 if the past information of 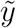 improves the prediction of the presence of 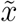 above and beyond what we could predict by the past of 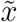 alone. That is, GC does not only assess the existence of a connection but also estimates the direction of that connection. We here use a spectrally resolved multivariate extension of GC (Geweke, 1982; Barrett et al., 2010; Barnett and Seth, 2014), which allows us to estimate Granger-causal influences between groups of variables at individual frequencies. There are multiple strategies to arrive at spectral Granger causality estimates. Here, we follow recommendations made in Barnett and Seth (2014, 2015); Faes et al. (2017); Barnett et al. (2018) that ensure stable and unbiased estimates, and use Matlab code provided by the respective authors.

We first transform the joint cross-spectrum into an autocovariance sequence **G**_[**xy**]_(*p*) ∈ ℝ^(*K*+*L*)×(*K*+*L*)^ with lags *p* ∈ {0, 1, …, *N*_*P*_ *}, N*_*P*_ = 20, using the inverse Fourier transform. The autocovariance spectrum is further used to estimate the parameters **A**(*p*) ∈ ℝ^(*K*+*L*)×(*K*+*L*)^, *p* ∈ {1, …, *N*_*P*_} and **Σ** = Cov_*t*_ [***ϵ***(*t*)] *∈* ℝ^(*K*+*L*)×(*K*+*L*)^ of a linear autoregressive model

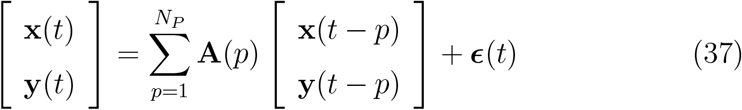

of order *N*_*P*_ using Whittle’s algorithm (Whittle, 1963; Barnett and Seth, 2014). Autoregressive model parameters are next converted into a statespace representation 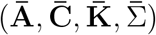 corresponding to the model

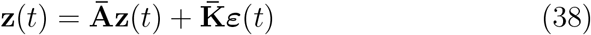

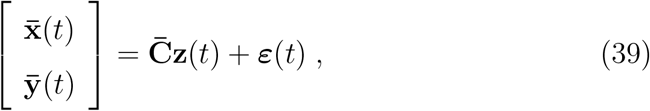

using the method of Aoki and Havenner (1991), where 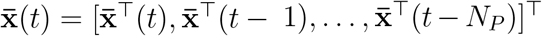 and 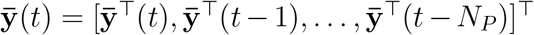 are temporal embeddings of order 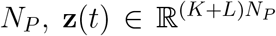 and 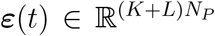 are unobserved variables, and all parameters are (*K* + *L*)*N*_*P*_ × (*K* + *L*)*N*_*P*_ matrices. Subsequently, the transfer function 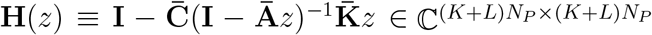 of a moving-average representation

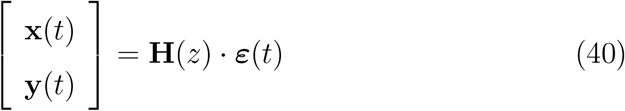

of the observations is derived, where 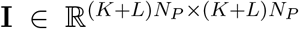 denotes the identity matrix and where *z* = *e*^*-i*4*πf/T*^ for a vector of frequencies *f ∈* {0 Hz, 0.5 Hz, …, 50 Hz}, *T* = 200, and a factorization of the joint crossspectrum is obtained as 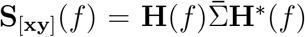 (Barnett and Seth, 2015).

Frequency-dependent *Granger scores*

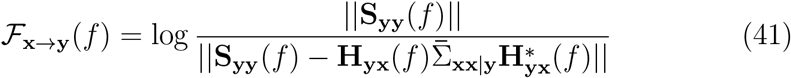

and (analogously) ℱ_**y**→**x**_(*f*) are then calculated, where **H**(*f*) and 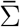 are partitioned in the same way as **S**(*f*), where 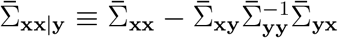 denotes a partial covariance matrix, and where || · || denotes matrix determinant (Barnett and Seth, 2015). Finally, differences

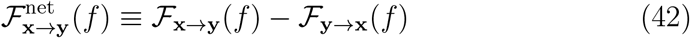

and 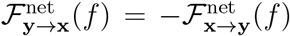 summarizing the net information flow between the multivariate time series 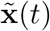 and 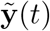 are calculated (Winkler et al., 2016).

Just like coherence, GC is not robust, i.e. can deliver spurious results for mixtures of independent sources as a result of volume conduction or source leakage (e.g., Haufe et al., 2012, 2013). This can be easily acknowledged by considering a single source that spreads into two measurement channels, which are superimposed by distinct noise terms. In that case, both channels will mutually improve each other’s prediction in the sense of GC (Haufe and Ewald, 2019). This problem is overcome by a robust version of GC, time-reversed GC (TRGC), which introduces a test on the temporal order of the time series. That is, TRGC estimates the directed information flow once on the original time series and once on a time-reversed version of the time series.

If GC is reduced or even reversed when the temporal order of the time series is reversed, it is likely that the effect is not an artifact coming from volume conduction (Haufe et al., 2012, 2013; Vinck et al., 2015; Winkler et al., 2016). Formally, multivariate spectral GC as introduced above can be evaluated on the time-reversed data by fitting the autoregressive model in Eq. (37) on the transposed autocovariance sequence 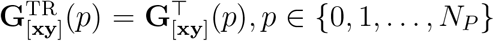.

This yields net GC scores 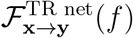 for the time-reversed data, which are subtracted from the net scores obtained for the original (forward) data to yield the final time-reversed GC scores:

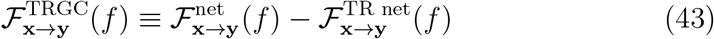

and (analogously)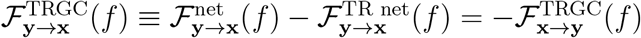

### 2.5. Pipelines

In the following section, we describe the processing pipelines that were tested. All pipelines take the sensor measurements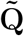 as input. Then all pipelines calculate and apply an inverse model **P** to project sensor data to source level. From there, we aggregate voxel activity within regions by employing PCA and estimate inter-regional FC with various FC metrics described above. We describe several strategies of combining PCA with the calculation of FC in the following subsections. This step results in a *N*_*roi*_ × *N*_*roi*_ × *N*_*freq*_ FC matrix which is then averaged across the frequency bins within the interaction frequency band (8-12 Hz). The output of all pipelines is one connectivity score for every region combination. We describe the processing exemplarily for the calculation of FC between two regions X and Y.

#### Pipelines FIXPC1 to FIXPC6: Fixed number of principal components

The first six pipelines use PCA dimensionality reduction. Afterwards, depending on the pipeline, a fixed number *C* of either one, two, three, four, five, or six strongest PCs are selected for further processing. Then, FC is calculated: in case of univariate measures (i.e., coherence and iCOH), we first calculate FC scores between all PC combinations of the two regions X and Y and then average across all pairwise FC scores. In case of multivariate FC measures, we directly calculate a single FC score between the PCs of region X and those of region Y. This approach has been used previously (e.g. Schoffelen et al., 2017).

#### Pipelines VARPC90 and VARPC99: Variable numbers of principal components

Pipelines VARPC90 and VARPC99 are equivalent to the FIXPC pipelines, with the difference that we do not select the same fixed number of PCs for every region. Instead, we select the number of PCs such that at least 90% (VARPC90) or 99% (VARPC99) of the variance in each ROI is preserved (c.f. Section 2.3). Thus, an individual number of PCs is chosen for each region. FC is then calculated analogously to pipelines FIXPC1 to FIXPC6. The idea of selecting the number of PCs such that a pre-defined fraction of the variance is retained has been used in previous literature (e.g. Gómez-Herrero et al., 2008).

#### Pipeline MEANFC: Mean first FC second

In this pipeline, the time series of all voxels within one region are averaged separately for the three orthogonal dipole orientations. Then, for univariate FC measures, FC is calculated between all 3*3 dimension combinations of the 3D-time series of region X and region Y. Afterwards, the average of these nine FC scores is taken. Multivariate FC measures are directly calculated between the 3D time series.

#### Pipeline CENTRAL: Central voxel pick

In this pipeline, we select only the central voxel of each region for further processing. The central voxel of a region is defined as the voxel whose average Euclidean distance to all other voxels in the region is minimal. To calculate the FC score between the 3D time series of the central voxel of region X and the 3D time series of the central voxel of region Y, we proceed analogous to pipeline MEANFC: in case of univariate FC measures, the FC score for all combinations of dipole orientations is calculated and then averaged. In case of multivariate FC measures, only one FC score is calculated between the two 3D time series. Selecting the time series of the central voxel as the representative time series for the region is an idea that has been used in previous studies already (Perinelli et al., 2022).

#### Pipeline FCMEAN: FC first mean second

In pipeline FCMEAN, the multivariate FC between each 3D voxel time series of region X with each voxel time series of region Y is calculated first. That is, if *R*_*X*_ is the number of voxels of region X and *R*_*Y*_ is the number of voxels in region Y, *R*_*X*_ * *R*_*Y*_ FC scores for all voxel combinations are calculated. To obtain a single FC score between region X and region Y, we then average all *R*_*X*_ * *R*_*Y*_ FC scores. Due to computational and time constraints, we test this pipeline only for MIM and MIC. This approach has also been used in the literature before (Babiloni et al., 2018).

#### Pipeline TRUEVOX: True voxel pick

This pipeline is used as a baseline. Here we select the voxel for further processing that indeed contains the activity of the given ROI—i.e. the ground-truth voxel (see Section 2.1). All further processing is analogous to pipeline CENTRAL. In configurations with two active voxels per region (see Section 3, Experiment 6), FC scores are calculated for 2 * 3 * 3 voxeland dipole orientation combinations.

### 2.6. Performance evaluation

We use a rank-based evaluation metric to assess the performance of the pipelines. All processing pipelines result in one FC score for every region– region combination. To evaluate the performance of a pipeline, we first sort all FC scores in a descending order and retrieve the rank 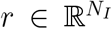, with *N*_*I*_ ∈ {1, 2, 3, 4, 5} denoting the number of ground-truth interactions. Based on this rank vector, we calculate the percentile rank (PR):

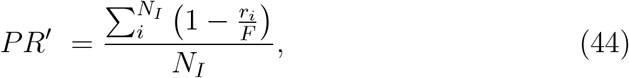

with *F* denoting the total number of FC scores. The *PR*′ is then normalized to the perfect-skill *PR*_*ps*_ and no-skill *PR*_*ns*_ cases, and is therefore defined between 0 and 1:

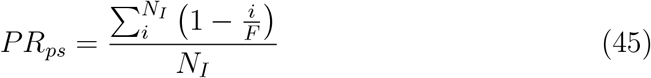

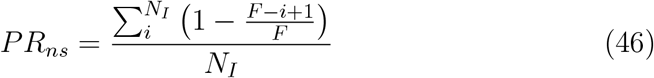

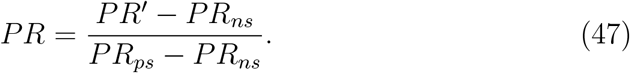

We report all PR values rounded to the second decimal. In case of the phase-based FC metrics, the PR is calculated on the original FC scores. In case of GC and TRGC, we separately evaluate each pipeline’s interaction detection ability, and its ability to determine the direction of the interaction. For evaluating the detection, we calculate the PR on the absolute values of the FC scores, whereas for evaluating the directionality determination performance, we calculate the PR only on the positive FC scores. Note that this is sufficient for the anti-symmetric directed FC measures used here.

### 2.7. Statistical assessment

In Experiment 1C, we provide a suggestion on how to statistically assess the presence of FC. Here, we obtain p-values by testing against a surrogate distribution consistent with the null hypothesis of zero interaction between all region pairs. The 10,000 samples of the surrogate distribution are drawn by shuffling epochs relative to each other when computing the cross-spectrum. More specifically, we calculate the cross-spectrum between the time series of one region and the shuffled time series of another region with the Welch method, where the diagonal entries of the cross-spectrum (spectral powers) are obtained without shuffling. From the shuffled cross-spectrum, MIM is calculated. We obtain p-values by counting the number of shuffled MIM-samples that are higher than the true MIM score and dividing this number by the total number of samples in the null distribution. FDR-correction (*α*-level = 0.05) is used on the upper triangle of the region–region p-value matrix to set a significance threshold.

### 2.8. ROIconnect toolbox

Based on our experimental results (see Section 3), we identified a set of recommended methods and pipelines. These have been implemented in a Matlab toolbox and are made available as a plugin to the free EEGlab package^1^. This toolbox also contains code for analyzing spectral power in EEG source space, and for visualizing power and FC results in source-space. A comprehensive description of the functionality and usage of the toolbox is provided in Appendix A. Moreover, an exemplary application of the toolbox to the analysis of a real EEG dataset is provided in Section 4.

## 3. Experiments and Results

We conducted a set of experiments to assess the influence of the different pipeline parameters on the reconstruction of ground-truth region-to-region FC. We describe the general experimental setting in Figure 2. Each experiment consisted of the following steps: (1) Signal generation. (2) Source projection. (3) Dimensionality reduction within regions. (4) Functional connectivity estimation. (5) Performance evaluation. Each experiment was carried out 100 times (= iterations). If not indicated otherwise, all experiments had the following default setting:

- LCMV inverse solution
- SNR = 3.5 dB
- BSR = 0 dB
- number of interactions = 2
- time delay of the interaction = 50 to 200 ms
- number of generated sources per region = 1

**Figure 2:**
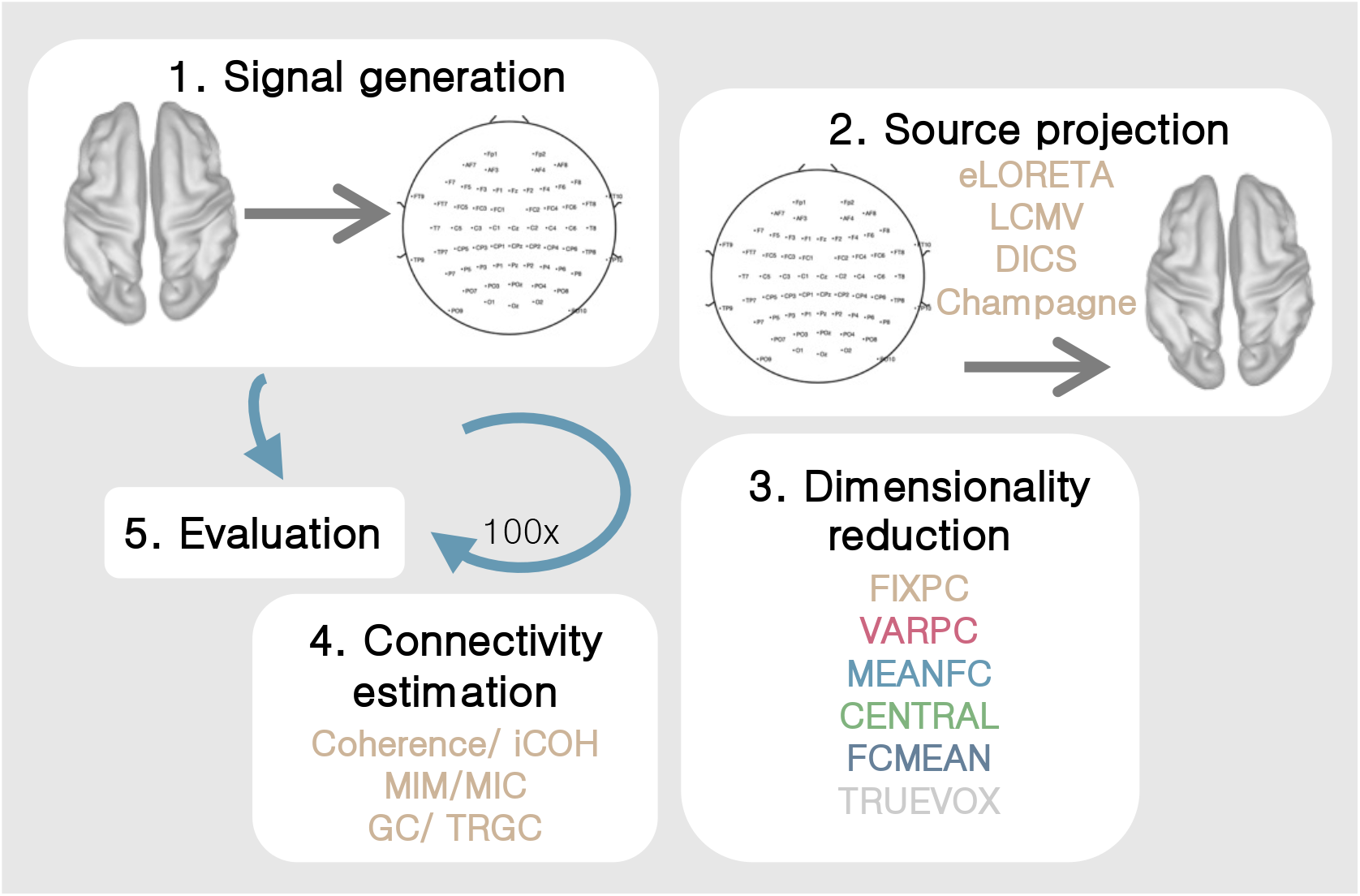
Experimental setup. Every experiment consisted of five consecutive steps: (1) Signal generation. (2) Source projection. (3) Dimensionality reduction within regions. (4) Functional connectivity estimation. (5) Performance evaluation. Every experiment was carried out 100 times.

If not stated otherwise, the following parameters were drawn randomly in each iteration: ground-truth interacting (seed and target) regions (two distinct regions uniformly drawn between 1 and *N*_*roi*_), ground-truth active voxel(s) within regions (uniformly drawn between 1 and *R*_roi_), time delay (uniformly drawn between 50 and 200 ms). Furthermore, brain noise and sensor noise, as well as the signal were generated based on (filtered) random white noise processes as described above.

Figure 3 to Figure 11 show the results of experiments 1–6. In addition, all main results are summarized in Table 1. All figures (plotting code adapted from Allen et al., 2019) follow the same scheme: in every subplot, the 100 dots on the right side mark the performance, i.e. the PR, measured in each of the 100 iterations. On the left, a smooth kernel estimate of the data density is shown. The red and black lines represent the mean and median PR of the experiment, respectively, and the boxcar marks the 2.5th and 97.5th percentiles. Please note that the Y-axis is scaled logarithmically in all plots. We tested differences between pipeline performances with a one-sided Wilcoxon signed-rank test. Please note that a p-value *p*_A,B_ corresponds to a one-sided test for *B > A*.

**Figure 3:**
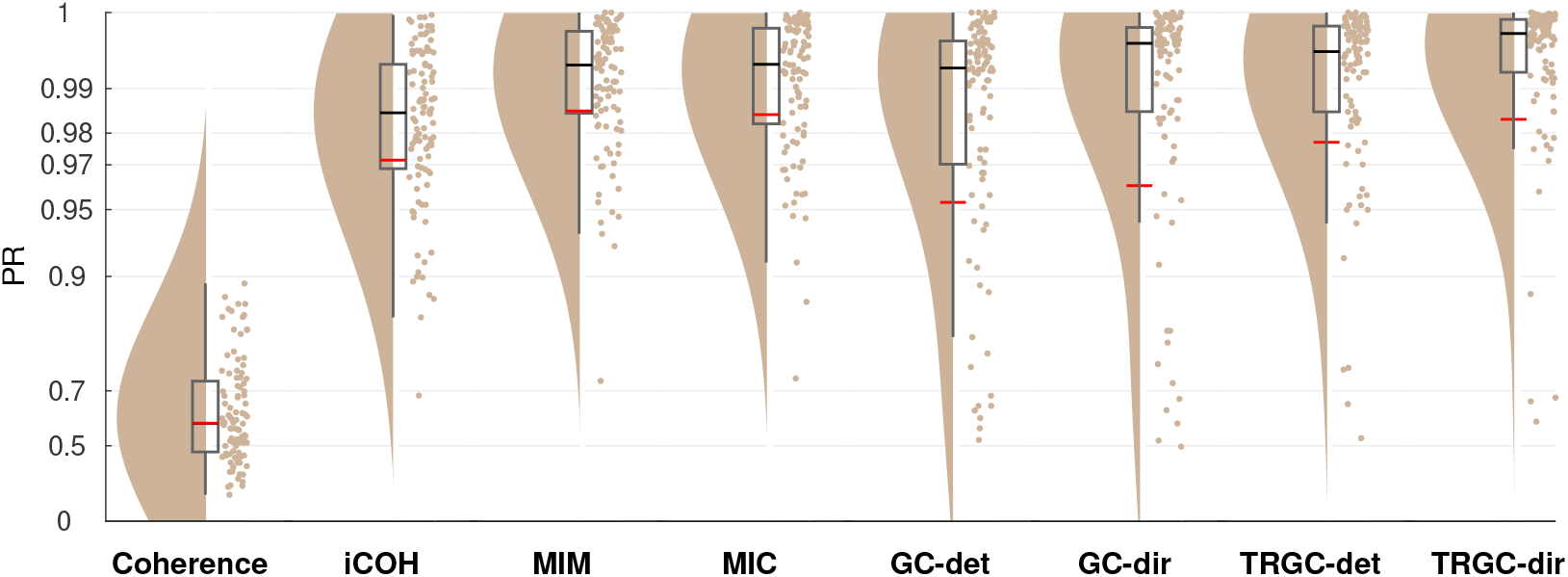
Comparison of different functional connectivity metrics (Experiment 1A). Red and black lines indicate the mean and median percentile rank (PR), respectively. The boxcar marks the 2.5th and 97.5th percentiles.

**Table 1:**
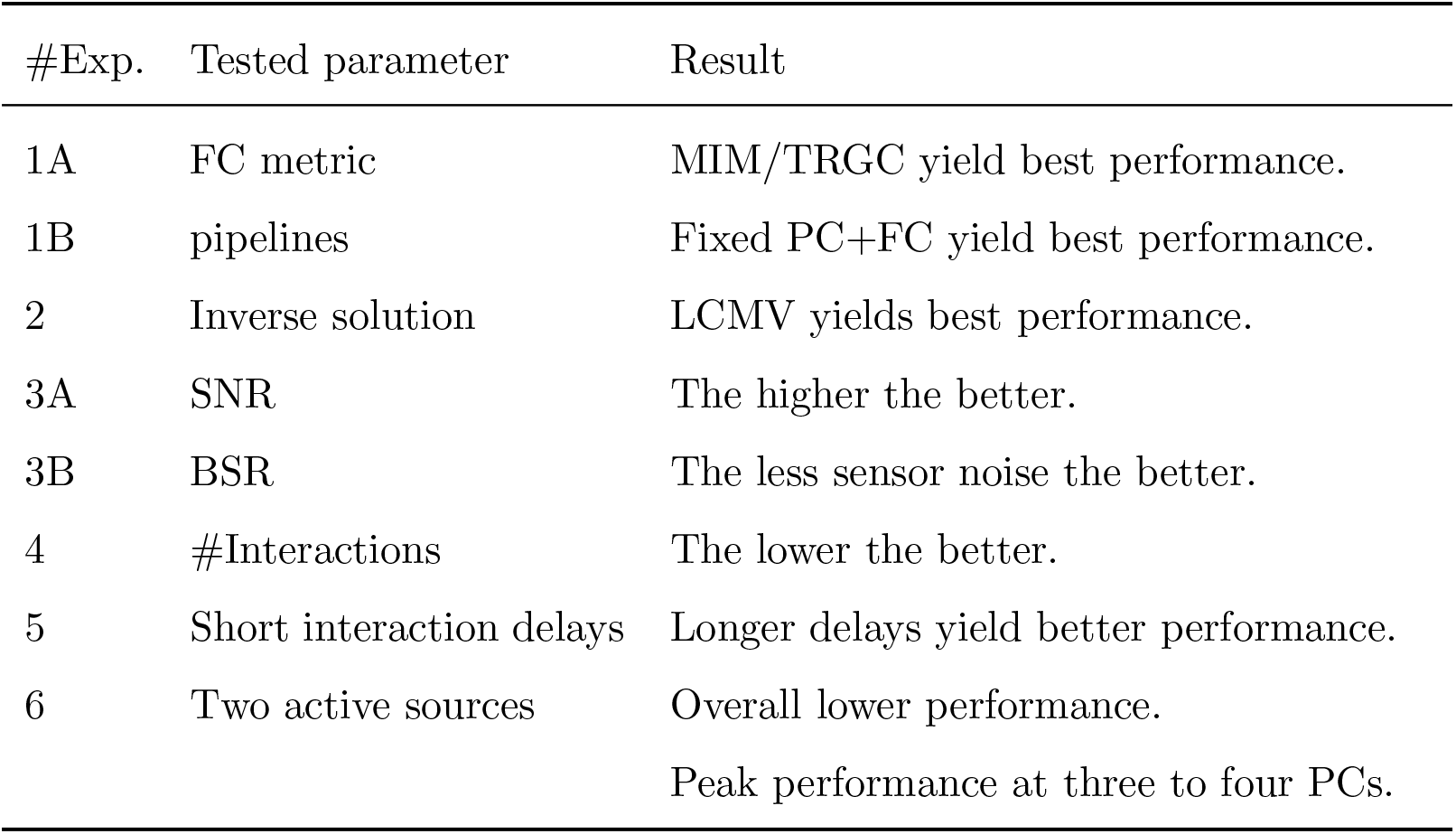
Summary of the results of experiment one to six. A pipeline including robust multivariate FC metrics like MIM or TRGC, a PCA with fixed number of selected components, and LCMV source reconstruction yields the best performance.

Matlab code to reproduce all experiments is provided under^2^.

### 3.1. Experiment 1

#### Experiment 1A

In Experiment 1A, we evaluated the performance of different FC metrics in detecting the ground-truth interactions. The ability to detect FC was tested for coherence, iCOH, MIC, MIM, GC, and TRGC. The ability to detect the correct direction of the interaction was tested for GC and TRGC (see Section 2.4).

In Figure 3, we show the performances of different FC metrics. We see that MIM, MIC and TRGC (detection) all have a mean PR of over 0.97 and clearly outperform the other measures in detecting the ground-truth FC. The non-robust metrics coherence (mean PR = 0.59) and GC (mean PR = 0.95) detect the ground-truth interactions less reliably (*p*_coherence, MIM_ < 10^−4^; *p*_GC,MIM_ = 0.0040). When comparing GC and TRGC in their ability to infer the direction of the interaction, TRGC (mean PR = 0.98) outperforms GC (mean PR = 0.96; *p*_GC,TRGC_ *<* 10^*-*4^).

#### Experiment 1B

In Experiment 1B, we tested the influence of different strategies of dimensionality reduction within regions. In Figure 4, we show the comparison for MIM (interaction detection) and TRGC (directionality determination). For MIM, we observe that the FIXPC pipelines show a better performance than most of the other pipelines. Within the FIXPC pipelines, the pipelines with two, three or four PCs perform best (all mean PR = 0.99, *p*_FIXPC5,FIXPC3_ < 10^−4^). Only the TRUEVOX (baseline) pipeline using ground-truth information on voxel locations expectantly shows a higher performance (mean PR = 1.00; *p*_FIXPC3,TRUEVOX_ < 10^−4^). The two VARPC pipelines show a substantially reduced performance (mean PR = 0.96 and mean PR = 0.73, respectively; both *p*_VARPC,FIXPC3_ < 10^−4^). The MEANFC and CENTRAL pipelines (mean PR = 0.98 and mean PR = 0.96, respectively) also show reduced performance in comparison to the FIXPC3 pipeline (both *p* < 10^−4^). The FCMEAN pipeline (mean PR = 0.97) also did not perform as well as the FIXPC3 pipeline (*p* < 10^−4^) while taking much longer to compute (FIXPC3 < 1 h, FCMEAN = 32 h, single core, allocated memory: 16 GB). In terms of directionality estimation using TRGC, the outcome is similar.

**Figure 4:**
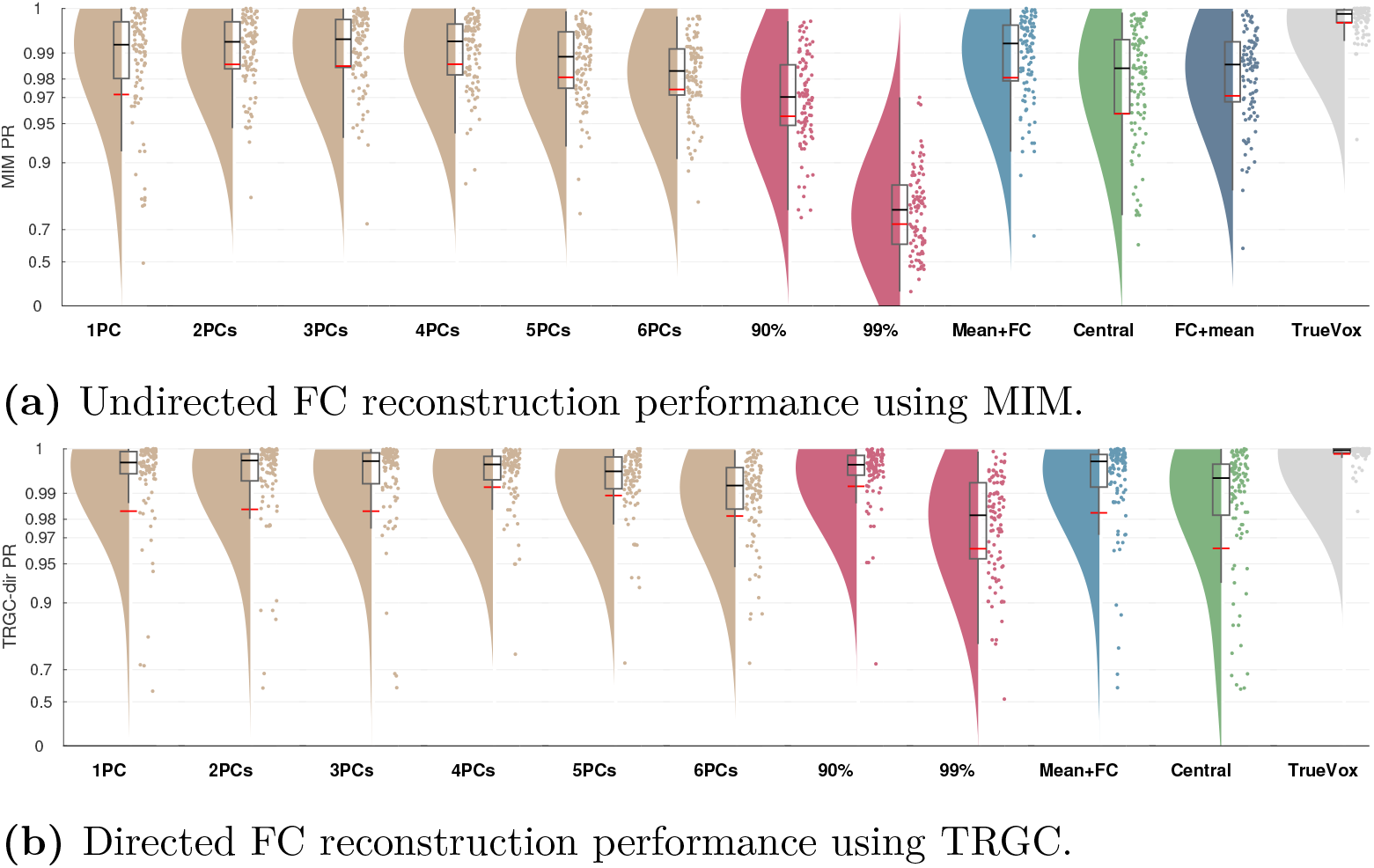
Comparison of different pipelines (Experiment 1B). (a) Undirected FC reconstruction performance achieved using the multivariate interaction measure (MIM). (b) Directed FC reconstruction performance achieved using time-reversed Granger causality. Red and black lines indicate the mean and median percentile rank (PR), respectively. The boxcar marks the 2.5th and 97.5th percentile.

Again, the TRUEVOX pipeline shows perfect performance (mean PR = 1.00). The FIXPC pipelines also exhibit very high performances (FIXPC4: mean PR = 0.99). Notably, in contrast to the results obtained with MIM, the VARPC90 also achieves competitive performance (mean PR = 0.99, *p*_VARPC90,FIXPC3_ = 0.0235). Please see Figure S1 to compare computation times of all pipelines.

We show the full matrix of all combinations of FC metrics and dimensionality reduction pipelines in Supplementary Figure S2. However, for all further experiments, we report performances only for MIM (interaction detection) and TRGC (directionality determination) since they performed best in Experiment 1A, and we focus on the FIXPC3 pipeline due the high performance observed in Experiment 1B.

#### Experiment 1C

To explore how to statistically assess the presence of FC, we performed an additional experiment for a specific setting (SNR = 3.5 dB, one interaction between region 11 and region 49, BSR = 0 dB, LCMV filter, dimensionality reduction to 3PCs, FC metric = MIM). Here, we obtained p-values by testing against a surrogate distribution consistent with the null hypothesis of zero interaction between all region pairs. In Figure 5, we contrast the ground-truth ROI-to-ROI connectome with the estimated FC per region combination as well as the -log10(p) values “surviving” the FDR-correction for this experiment. While in the ground-truth connectome only the ground-truth region combination shows a high MIM score, there are also some high MIM scores in other region combinations than the ground truth in the reconstructed source-level connectome. Still, the ground-truth region combination in this setting still achieves the second-highest MIM score (PR = 0.9996). However, in Figure 5c, we see that testing the statistical significance with a shuffling test results in a substantial number of significant false positive interactions in the vicinity of the simulated interacting region pair. We discuss this result in Section 5.

**Figure 5:**
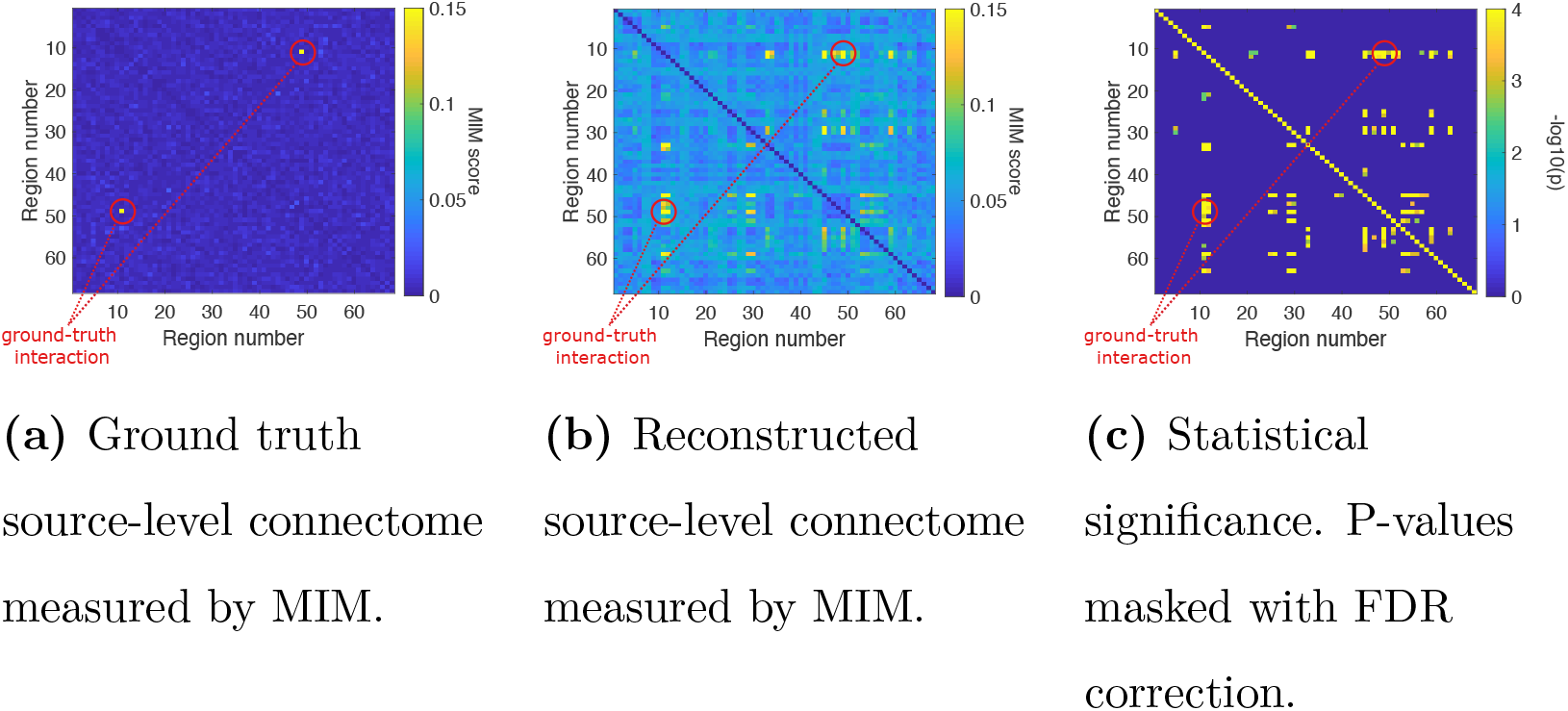
Comparison of the ground-truth ROI-to-ROI connectome with the estimated functional connectivity per region combination and the -log10(p) values after FDR-correction for a single experiment. A ground-truth interaction is modeled between region 11 and region 49.

### 3.2. Experiment 2

#### Experiment 2A

In Experiment 2, we tested the influence of the type of inverse solution on the pipelines’ performances. In Figure 6, we show the comparison between eLORETA, LCMV, DICS, and Champagne. We observe that the two beamformer solutions and Champagne clearly outperform eLORETA (mean PR 0.65; Figure 6a) in detecting undirected connectivity (all *p* < 10^*-*4^). While DICS, LCMV and Champagne all show very good performances, we see a slight advantage of LCMV (mean PR = 0.99) in comparison to Champagne (mean PR = 0.97, *p*_Champagne,LCMV_ = 0.0013). We do not observe a significant difference between DICS and LCMV (*p*_DICS,LCMV_ = 0.2805).

**Figure 6:**
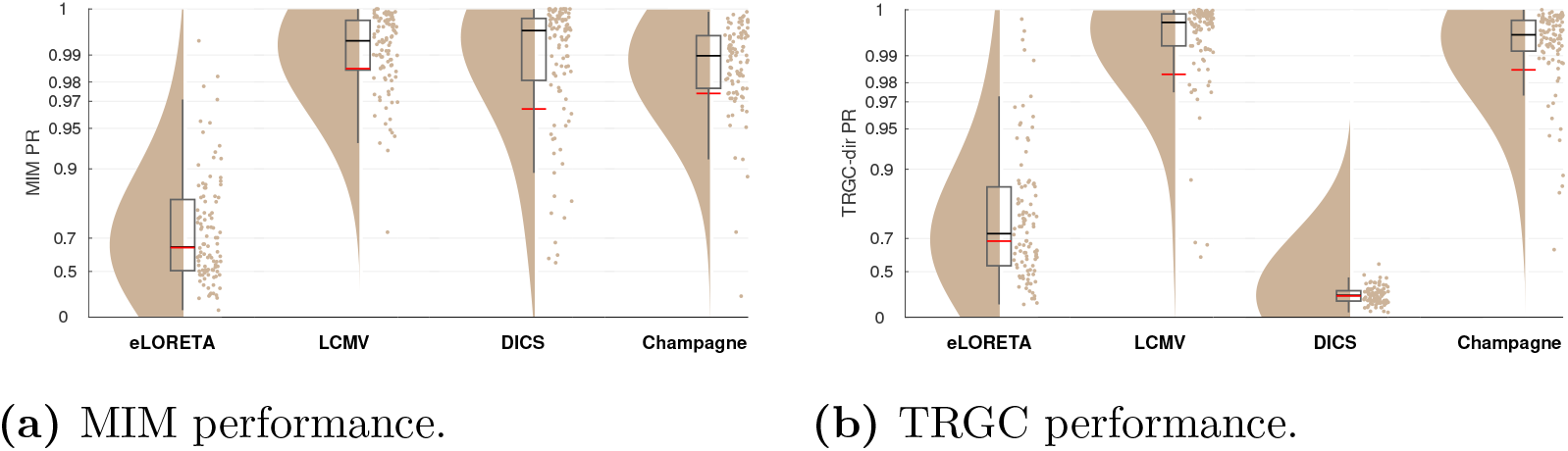
Comparison of different inverse solutions (Experiment 2). (a) Undirected FC reconstruction performance achieved using the multivariate interaction measure (MIM). (b) Directed FC reconstruction performance achieved using time-reversed Granger causality. Red and black lines indicate the mean and median percentile rank (PR), respectively. The boxcar marks the 2.5th and 97.5th percentile.

In terms of directionality determination (Figure 6b), the picture is different: while LCMV (mean PR = 0.98) leads to accurate directionality estimates, DICS fails to detect the direction of the ground-truth interaction in a high number of experiments (mean PR = 0.28, *p*_DICS,LCMV_ < 10^−4^). eLORETA also shows a reduced overall performance (mean PR = 0.69, *p*_eLORETA, LCMV_ < 10^−4^). Champagne shows decent performance (mean PR = 0.99), which is, however, lower than that of LCMV (*p*_Champagne,LCMV_ < 10^−4^).

The differences in computation times of the different inverse solutions are also remarkable. While LCMV (2 sec) and DICS (178 sec) are fast to compute, eLORETA (388 sec) and Champagne (3747 sec) take much longer to compute as a cross-validation scheme to set the regularization parameter is implemented for both. Setting the regularization parameter to a default value would drastically reduce computation time for eLORETA and Champagne, but would also decrease performance (results not shown).

#### Experiment 2B

To investigate further why eLORETA performs considerably less well than LCMV in our experiments, we generated ground-truth activity with an interaction between one seed voxel in the left frontal cortex and one target voxel in the left precentral cortex. We then again generated sensor data as described in Section 2.1 and applied pipeline FIXPC1 to calculate regional MIM scores. In Supplementary Figure S3, we show the resulting power maps, as well as seed MIM scores and target MIM scores for data projected with eLORETA and MIM, respectively. We see clearly the advantage of LCMV: while both power and MIM in the eLORETA condition are spread out to other regions, LCMV is able to localize the ground-truth power and connectivity very precisely.

#### Experiment 2C

Does LCMV only perform so well in our experiment because our experimental setup artificially favors it? In the following additional analysis, we investigated whether LCMV still has an advantage over eLORETA when multiple pairs of correlated sources are present. More specifically, we here simulated two pairs of interacting sources where the time courses of the second source pair were identical to those of the first source pair. Results are presented in Figure 7. Please note that in this case, also the cross-interactions between the seed and target regions were evaluated as ground-truth interactions. We see that, while eLORETA is not much affected by the correlated sources setup, LCMV has a decreased reconstruction performance according to both MIM and TRGC. However, LCMV still performs better than eLORETA even in this setup (*p*_eLORETA,LCMV_ < 10^*-*4^).

**Figure 7:**
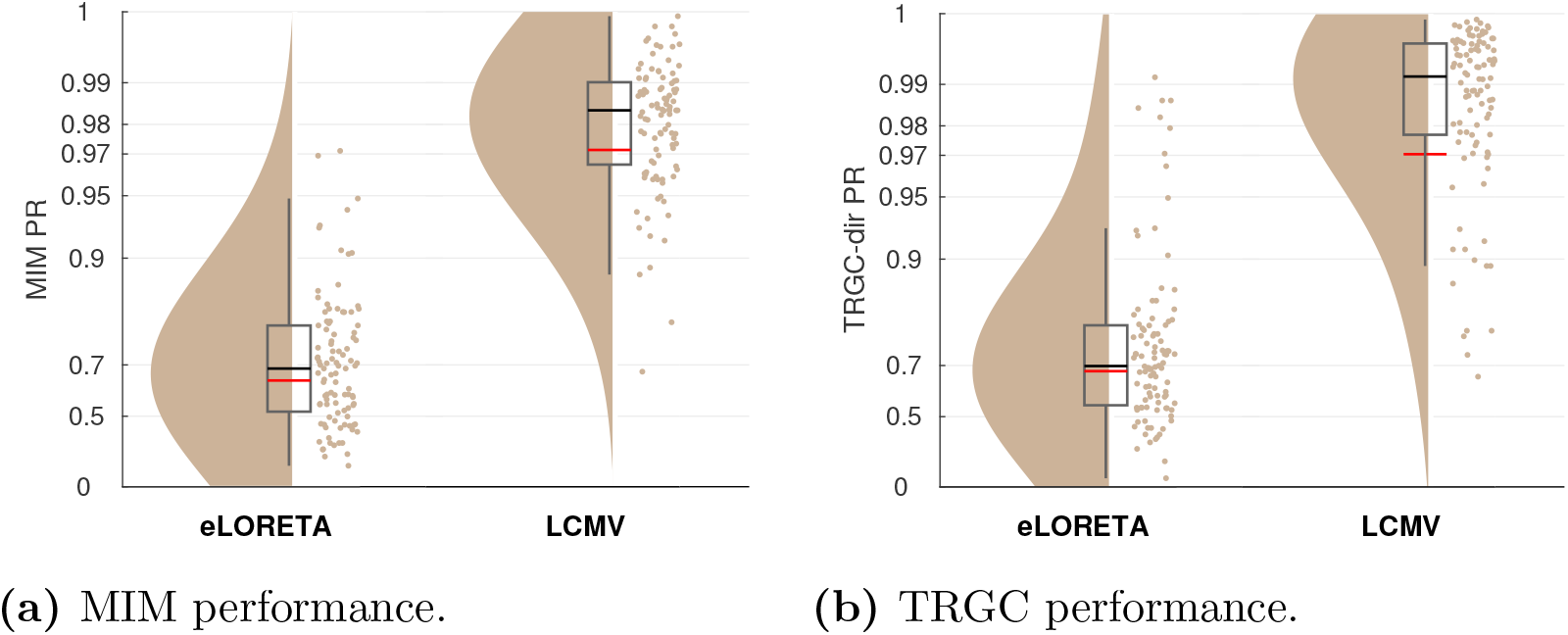
Performance observed for two perfectly correlated source pairs. (a) Undirected FC reconstruction performance achieved using the multivariate interaction measure (MIM). (b) Directed FC reconstruction performance achieved using time-reversed Granger causality. Red and black lines indicate the mean and median, respectively. The boxcar marks the 2.5th and 97.5th percentile.

#### 3.3. Experiment 3

In real-world EEG measurements, data are to a certain extent corrupted by noise, e.g. from irrelevant brain sources, or by noise sources from the outside. In Experiment 3, we investigated the effect of SNR and BSR on FC estimation performance. In Figure 8a and 8b, we show the performance of the FIXPC3 pipeline for SNRs of -7.4 dB, 3.5 dB and 19.1 dB. For both MIM (Figure 8a) and TRGC (Figure 8b), we observe decreased performances for decreased SNRs, as expected. For an SNR of 19.1 dB, nearly all experiments show a perfect detection of ground-truth interactions (mean PR > 0.99).

**Figure 8:**
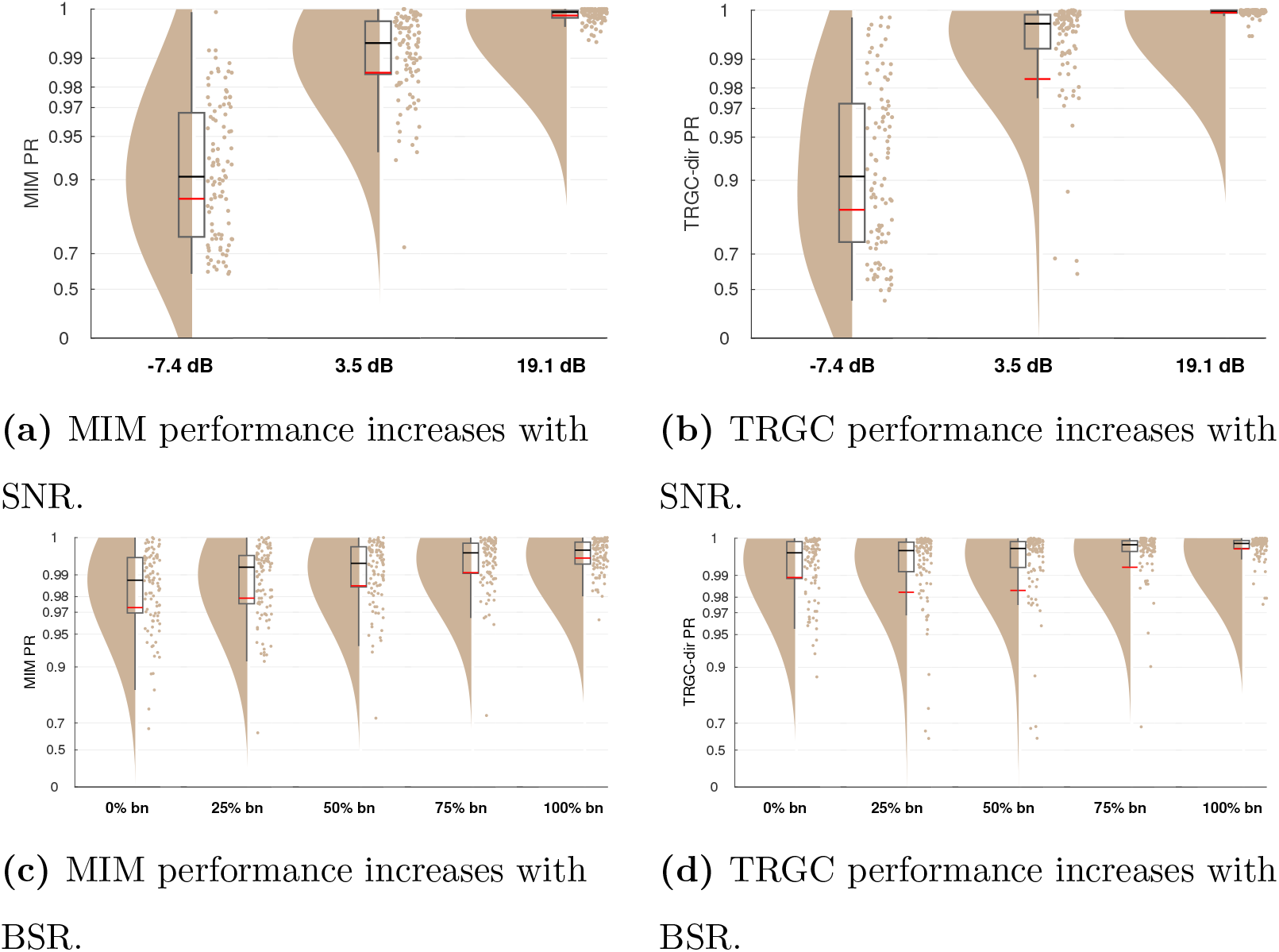
FC estimation performance depends on the signal-to-noise ratio and brain noise-to-sensor noise ratio (Experiment 3). (a/c) Undirected FC reconstruction performance achieved using the multivariate interaction measure (MIM). (b/d) Directed FC reconstruction performance achieved using time-reversed Granger causality. Red and black lines indicate the mean and median percentile rank (PR), respectively. The boxcar marks the 2.5th and 97.5th percentile.

Is FC detection more impaired by pink brain noise or white sensor noise? In Experiment 3B, we tested the performance for BSR environments of 100% sensor noise, 25% brain noise, 50 % brain noise, 75% brain noise, and 100% brain noise. In Figure 8c and 8d, we show the performances for different BSRs. We observe a slightly better performance for signals more strongly contaminated by correlated brain noise than white sensor noise (mean MIM PR 100% brain noise > 0.99) compared to the opposite case (mean MIM PR 0% brain noise = 0.97).

Note that in Experiments 1 to 3, for better comparison between the experimental conditions and to avoid variation due to random factors besides the experimental variation, we used the same generated data within an iteration in every experiment and only varied the tested condition.

### 3.4 Experiment 4

While we focused on a very simple scenario with only two interacting region pairs so far, real brain activity likely involves multiple interacting sources. To increase the complexity in our setup, we compared performances for different numbers of interacting region pairs in Experiment 4. As expected, Figure 9 clearly shows that more simultaneous true interactions lead to decreased ability to reliably detect them. While the detection is nearly perfect for one interaction (mean MIM PR > 0.99; mean TRGC PR > 0.99), the performance is much reduced for 5 interactions (mean MIM PR = 0.91; mean TRGC PR = 0.93). This applies for both MIM and TRGC. Please note however, that despite using a normalized version of the PR (see Section 2.6), the PR metric is not perfectly comparable for different numbers of true interactions. That is, when calculating the PR on randomly drawn data, the PR distribution is close to uniform when only one interaction is assumed, but shows a normal distribution with increasing kurtosis for higher numbers of interactions. However, the mean of the distribution equals to 0.5 for all assumed interactions.

**Figure 9:**
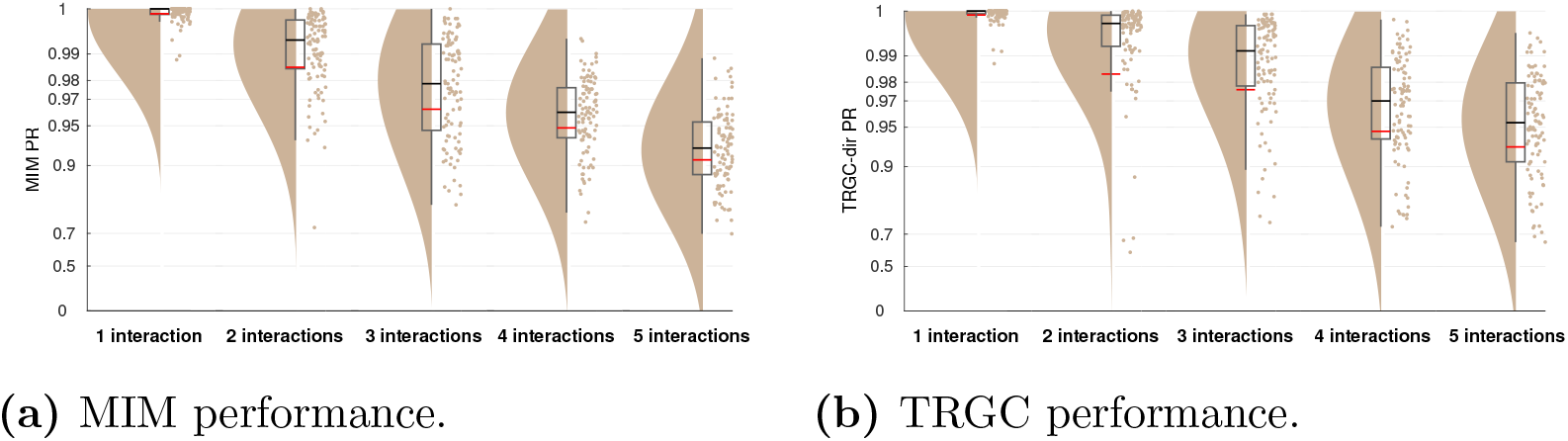
FC reconstruction performance depends on the number of true interactions (Experiment 4). (a) Undirected FC reconstruction performance achieved using the multivariate interaction measure (MIM). (b) Directed FC reconstruction performance achieved using time-reversed Granger causality. Red and black lines indicate the mean and median percentile rank (PR), respectively. The boxcar marks the 2.5th and 97.5th percentile.

### 3.5. Experiment 5

While it is not entirely clear how large interaction delays in the brain can be, they likely range between 2 and 100 ms, depending not only on physical wiring, but also on cognitive factors (see Section 5). In Experiment 5, we evaluated to which degree the performance drops when regions interact with shorter time delays of 2, 4, 6, 8, and 10 ms. While the performance for the MIM metric is already quite impaired for a delay of 10 ms (mean PR = 0.90), performance drops drastically for 4 ms (mean PR = 0.73) and 2 ms (mean PR = 0.60) (Figure 10a). Detecting the direction of the interaction with TRGC is already much more difficult at a true delay of 10 ms (mean PR = 0.73) and is further reduced for a delay of 2 ms (mean PR = 0.56; Figure 10b).

**Figure 10:**
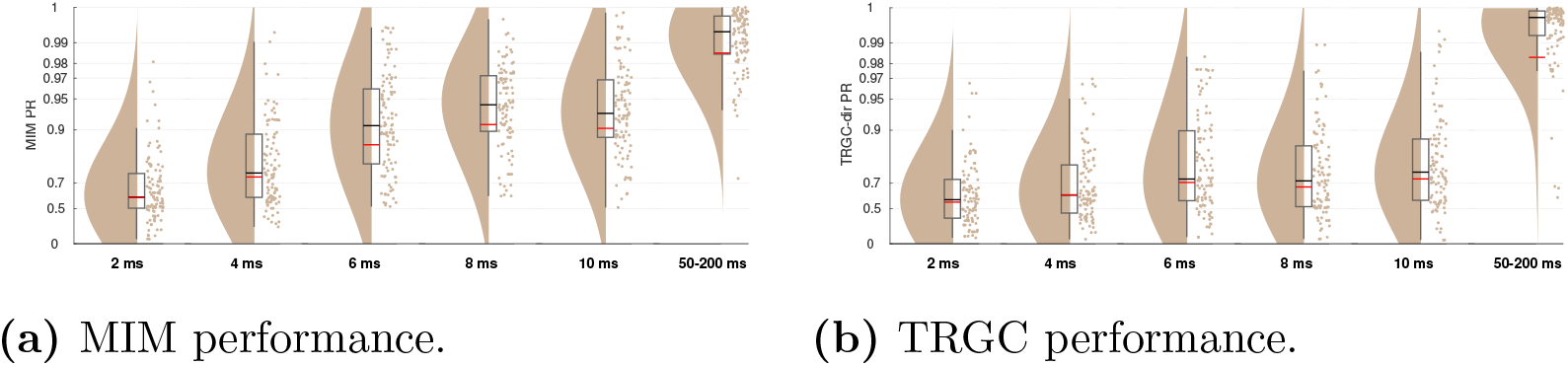
Performance for very small interaction delays and the default delay (Experiment 5). (a) Undirected FC reconstruction performance achieved using the multivariate interaction measure (MIM). (b) Directed FC reconstruction performance achieved using time-reversed Granger causality. Red and black lines indicate the mean and median percentile rank (PR), respectively. The boxcar marks the 2.5th and 97.5th percentile.

### 3.6. Experiment 6

In our previous experiments, the FIXPC pipelines with two to four PCs showed the best performance. But the ‘optimal’ number of PCs likely depends on the number of (interacting and non-interacting) signals in the brain as well as their relative strengths. To verify that the optimal number of PCs depends on the number of true sources, we increased the number of active voxels per region to two in Experiment 6. We then simulated two bivariate interactions between two different source pairs originating from the same regions.We show the results for pipelines FIXPC1 to FIXPC6. Interestingly, we here see that pipelines FIXPC3 (mean MIM PR = 0.99; mean TRGC PR = 0.99) and FIXPC4 (mean MIM PR = 0.99; mean TRGC PR = 0.99) perform clearly better than FIXPC1 (mean MIM PR = 0.89; mean TRGC PR = 0.93) or FIXPC6 (mean MIM PR = 0.98; mean TRGC PR = 0.98).

Based on these results, we confirm that the choice of the optimal number of fixed PCs increases with the number of independently active processes within one region (see Section 5 for further discussion).

## 4. Exploratory analysis of functional connectivity in left vs right motor imagery

To illustrate how the recommended analysis pipeline can be used to analyse real EEG data, we show an exploratory analysis of power and FC in left and right motor imagery. In the Berlin arm of the so-called VitalBCI study (Blankertz et al., 2010; Sannelli et al., 2019), 39 subjects conducted an experiment in which they imagined a movement with either the left or the right hand (Motor Imagery Calibration set; MI-Cb 1-3). Each trial consisted of a visual stimulus showing a fixation cross imposed with an arrow indicating the task for the trial (i.e., left or right motor imagery). After 4 sec, the stimulus disappeared, and the screen stayed black for 2 sec. Every subject conducted 75 left and 75 right motor imagery trials. During the experiment, EEG data were recorded with a 119-channel whole-head EEG system with a sampling rate of 1000 Hz. For this study, we used a 90-channel whole head standard subset of them. For our analysis, we selected only the 26 subjects for which previous studies have reported that the left vs. right motor imagery conditions could be well separated using statistical and machine learning techniques (‘Category I’ in Sannelli et al., 2019). Further experimental details are provided in Blankertz et al. (2010); Sannelli et al. (2019).

We filtered the data (1 Hz high-pass filter, 48-52 Hz notch filter, and 45 Hz low-pass filter, all zero-phase forward and reverse second-order digital high-pass Butterworth filters), and then sub-sampled them to 100 Hz. We then rejected artifactual channels based on visual inspection of the power spectrum and the topographical distribution of alpha power (between zero and five per participant, mean 1.19 channels) and interpolated them (spherical scalp spline interpolation). A leadfield was computed using the template head model Colin27 5003 Standard-10-5-Cap339 that is already part of the EEGLAB toolbox. We then epoched the data from 1 to 3 sec post-stimulus presentation start and separated left from right motor imagery trials.

We used the `pop_roi_activity` function of the newly developed ROIconnect plugin for EEGLAB to calculate an LCMV source projection filter, apply it to the sensor data, and calculate region-wise power (see Appendix A for a more detailed description). We then normalized the power with respect to the total power between 3 and 7 Hz as well as 15 and 40 Hz, and averaged it across frequencies between 8 and 13 Hz. The statistical significance of the differences between right and left hand motor imagery power was assessed with a paired t-test in every region. In Figure S4, we show the negative log10-transformed p-values, multiplied with the sign of the t-statistic. As expected, the results show a clear lateralization for the activation of the motor areas.

To estimate inter-regional FC, we used the `pop_roi_connect` function to calculate MIM based on the three strongest PCs of every region. Again, MIM was averaged across frequencies between 8 and 13 Hz. To reduce the region-by-region MIM matrix to a vector of net MIM scores, we summed up all MIM estimates across one region dimension.

Analogous to our statistical evaluation of simulated data, described in Experiment 1C, we assessed the statistical significance of the net FC of each region against the null hypothesis of zero net interaction separately for each of the two motor imagery conditions. Specifically, we first calculated the true MIM score between all region pairs in all subjects. Then, we generated a null distribution of 1000 shuffled MIM scores for every region combination in every subject. Subsequently, the true and shuffled net MIM scores were calculated by averaging across one of the region dimensions. To obtain p-values, we compared the true MIM of every region and subject to the respective null distribution. To aggregate the p-values across subjects, we applied Stouffer’s method (see, e.g., Dowding and Haufe, 2018). Finally, FDR-correction (*α*-level = 0.05) was used to correct for multiple comparisons. We show the negative log10-transformed p-values in Figures 12a and 12b.

Additionally, we assessed the statistical difference between the net MIM scores of the left vs. right hand motor imagery condition by again using a paired t-test for every region. In Figure 12c, we show the negative log10-transformed p-values, multiplied with the sign of the t-statistic. Again, as expected, the results show a lateralization for the undirected net FC of the motor areas. Matlab code of the analyses presented in this section is provided under ^3^

## 5. Discussion

Estimating functional connectivity between brain regions from reconstructed EEG sources is a promising research area that has generated a number of important results (e.g. Hipp et al., 2011; Schoffelen et al., 2017; Babiloni et al., 2018). However, respective analysis pipelines consist of a number of subsequent steps for which multiple modeling choices exist and can typically be justified. In order to identify accurate and reliable analysis pipelines, simulation studies with ground-truth data can be highly informative. However, most existing simulation studies do not evaluate complete pipelines but focus on single steps. In particular, various published studies assume the locations of the interacting sources to be known a-priori, while, in practice, they have to be estimated as well. To this end, it has become widespread to aggregate voxel-level source activity within regions of an atlas before conducting FC analyses across regions. Multiple ways to conduct this dimensionality reduction step have been proposed, which have not yet been systematically compared using simulations. The main focus of our study was thus to identify those EEG processing pipelines from a set of common approaches that can detect ground-truth inter-regional FC most accurately. For the scenario modelled in this study, we observe that a pipeline consisting of an LCMV source projection, PCA dimensionality reduction, the selection of a fixed number of principal components for each ROI, and a robust FC metric like MIM or TRGC results in the most reliable detection of groundtruth FC (see Table 1). Consistent with results reported in Anzolin et al. (2019), LCMV consistently yielded higher FC reconstruction performance than eLORETA. Thus, we here answer the question that Mahjoory et al. (2017) left open, namely which source reconstruction technique is most suitable for EEG FC estimation. Our results are also in line with a larger body of studies that highlighted the advantages of robust FC metrics compared to non-robust ones (e.g. Nolte et al., 2004; Haufe et al., 2013; Vinck et al., 2015; Winkler et al., 2016; Schoffelen and Gross, 2019).

### Inverse solutions

For some inverse solutions, the choice of the regularization parameter has been shown to influence the accuracy of source reconstruction (Hincapié et al., 2016; Hashemi et al., 2021). While the parameter is of little importance for methods like LCMV and DICS, which are fitted separately to each source and thus solve low-dimensional optimization problems, it should be carefully chosen for full inverse solutions like Champagne and eLORETA, which estimate the activity at each source voxel within a single model. To avoid a performance drop due to unsuitable regularization parameter choice in eLORETA and Champagne, we used the spatial cross-validation method described in (Habermehl et al., 2014; Hashemi et al., 2021). This method automatically sets the parameter based on the data at hand and has been shown to improve the source reconstruction (Hashemi et al., 2021).

As hypothesized, DICS resulted in poor directionality determination performance, while LCMV and TRGC performed well. This can be explained by the difference between LCMV and DICS: while LCMV estimates the inverse solution in the time domain, DICS estimates the source projection for every frequency separately (Gross et al., 2001). This can lead to inconsistencies across frequencies. Since directionality estimation requires the aggregation of phase information across multiple frequencies, such inconsistencies may lead to failure of detecting true interactions and their directionalities. Therefore, we recommend to avoid using DICS source reconstruction when analysing directed FC. For undirected FC measures, this seems to be less of a problem. Still, in our simulation, LCMV consistently performed (even if only slightly) better than DICS. This can be explained by the lower effective number of data samples that are available to DICS at each individual frequency compared to LCMV, which uses data from the entire frequency spectrum. However, there may be cases when using DICS could result in more accurate localization. For example, this could be the case when the noise has a dominant frequency that is different from the signal.

### Robust functional connectivity metrics

In this study, we observed a strong benefit of using robust FC metrics over non-robust metrics in detecting genuine neuronal interactions. Overall, the performance of coherence is highly impaired by the volume conduction effect (see Figure 3, c.f. Nolte et al., 2004). The TRGC metric performed well for the investigation of the interaction direction, but also satisfyingly well for the interaction detection. However, the computation time for calculating TRGC exceeds that of MIM by far. Thus, we recommend using MIM to detect undirected FC in case the direction of the effect is not of relevance. If TRGC is calculated for estimating the direction of interactions, the absolute value of TRGC can be used to detect interactions as well.

Interestingly, GC without time reversal did not perform much worse than TRGC. This is in line with previous results (Winkler et al., 2016) demonstrating that the calculation of net GC values already provides a certain robustification against volume conduction artifacts. Concretely, it has been shown that net GC is more robust to mixed noise than the standard GC; however not as robust as TRGC (Winkler et al., 2016). We generally recommend using robust FC connectivity metrics like iCOH, MIM/MIC, or TRGC.

### Aggregation within regions

When comparing different processing pipelines, we found that employing an SVD/PCA and selecting a fixed number of components for further processing performs better than selecting a variable number of components in every ROI. When further investigating this effect, we found that, for MIM and MIC, the final connectivity score of the VARPC pipelines was positively correlated with the number of voxels of the two concerning ROIs (90%: MIM: *r* = 0.50, MIC: *r* = 0.32; 99%: MIM: *r* = 0.70, MIC: *r* = 0.41). This indicates that the flexible number of PCs leads to a bias in MIM and MIC depending on the size of the two involved ROIs. This could be expected, as the degrees of freedom for fitting MIM and MIC scale linearly with the number of voxels within a pair of regions. These in- or explicit model parameters can be tuned to maximize the FC of the projected data, which may lead to over-fitting. For finite data, this leads to a systematic overestimation of FC, to the degree of which it correlates with the number of voxels. Although representing a multivariate technique as well, similar behavior was not observed for TRGC. Here it is likely that a potential bias of the signal dimensionalities would cancel out when taking differences between the two interaction directions as well as between original and time-reversed data.

An interesting and so far unsolved question is how many fixed components should be chosen for further processing. In Experiment 6, we observed a clear performance peak around three to four components (Figure 11). In the default version with only one active source per ROI, we saw a similar pattern, but not as pronounced as in Experiment 6. This points towards a datadependent optimal number of components. Future work should investigate how this parameter can be optimized based on the data at hand.

**Figure 11:**
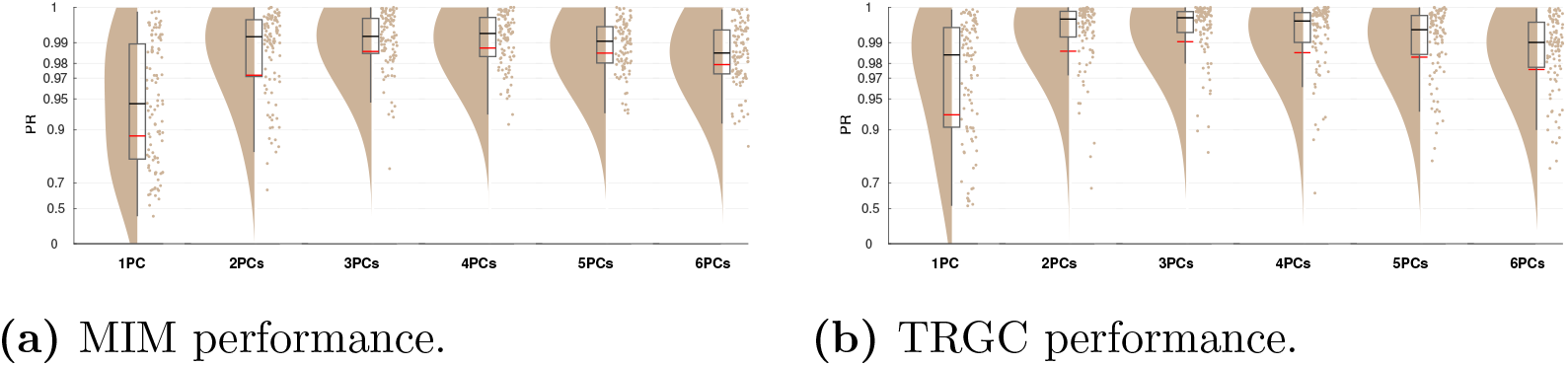
Performance when two active sources per region are simulated (Experiment 6). (a) Undirected FC reconstruction performance achieved using the multivariate interaction measure (MIM). (b) Directed FC reconstruction performance achieved using time-reversed Granger causality. Red and black lines indicate the mean and median, respectively. The boxcar marks the 2.5th and 97.5th percentile.

**Figure 12:**
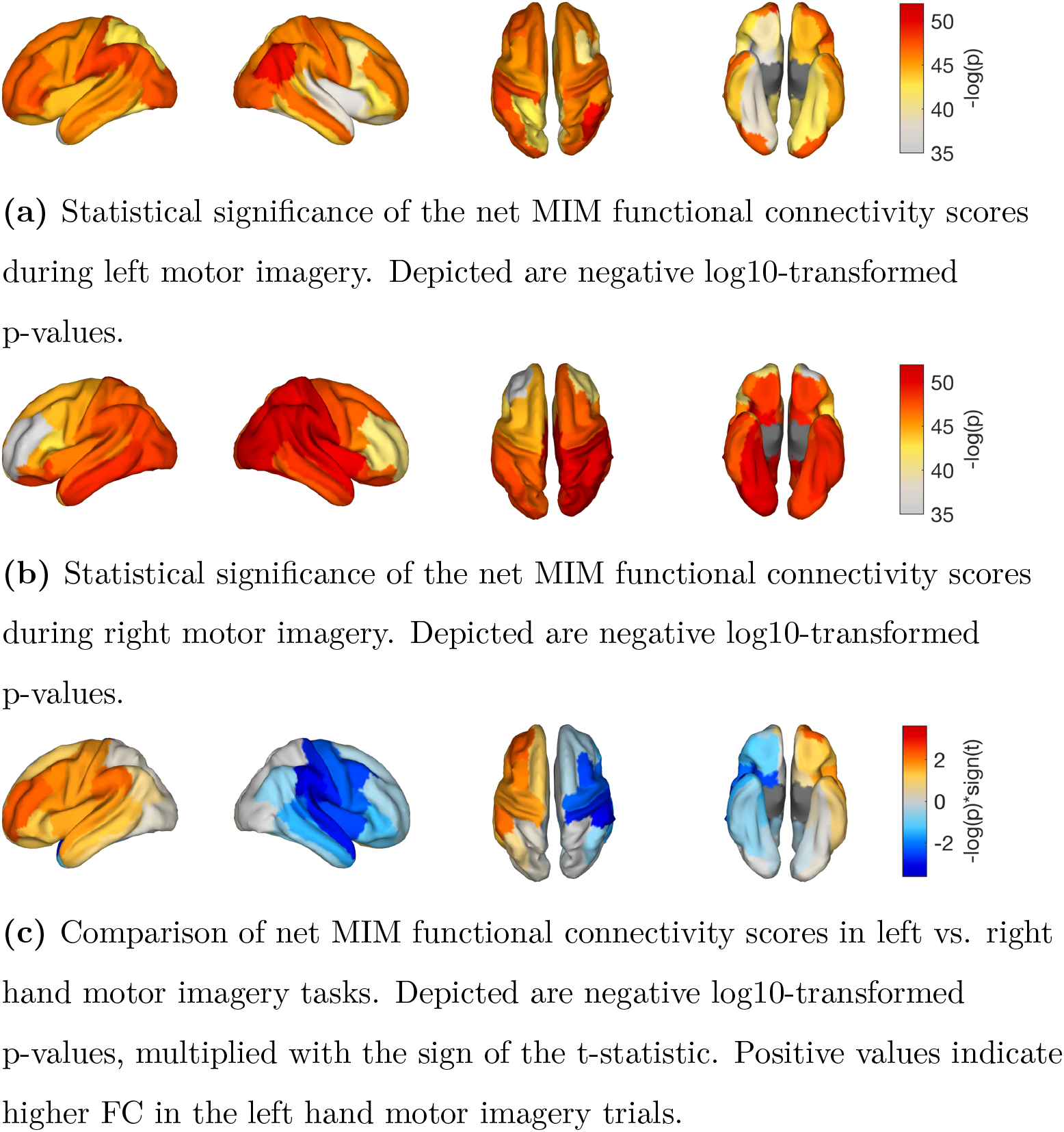
Results of the exploratory analysis of functional connectivity in left and right hand motor imagery tasks.

### Short time delays

In Experiment 5, we investigated to what extent the performance drops when the true interaction occurs with a very small time delay of 2 to 10 msec, which might be a realistic range for a number of neural interaction phenomena in the brain. Precise data on the typical order of the times within which macroscopic neural ensembles exchange information are, however, hard to obtain, as these transmission times depend not only on the physical wiring but also on cognitive factors that are not straightforward to model. Previous work has shown that delays can range from 2 to 100 msec, depending on the distance and number of synapses between two nodes (e.g. Fries, 2005; Oswal et al., 2016; Shouno et al., 2017; Miocinovic et al., 2018). For example, Oswal et al. (2016) studied interaction delays between the subthalamic nucleus and the motor cortex and found interaction delays of 20 to 46 msec. The satisfactory performance observed in our study for undirected FC at delays of 8 and 10 msec may therefore be of particular importance for clinical scientists that aim at investigating such long-range interactions. Note that the range of delays that can be detected with robust connectivity metrics strongly depends on the frequency band in which the interaction takes place. If the delay is very short compared to the base frequency of the interaction, then the phase difference it induces is close to either 0 or *±π*, making it less and less distinguishable from a pure volume conduction effect as it approaches these limits. In addition, the directionality of an interaction can only be resolved by analyzing multiple frequencies. Here, wider interaction bands lead to better reconstructions of the directionality of interactions with shorter delays, whereas higher frequency resolutions (that is, longer data segments) lead to better reconstructions of the directionality of interactions with longer delays. Here, we have demonstrated that alpha-band interactions with physiologically plausible transmission delays can be detected at 0.5 Hz frequency resolution, depending on the underlying SNR as well as additional modeling assumptions (see Limitations below).

### Statistical assessment

The goal of this study was to evaluate data analysis pipelines to assess FC. However, we excluded the assessment of any subsequent statistical evaluation of FC, which is not straightforward to investigate in simulation studies. In a simulation setting, we are free to choose the two factors that influence the statistical power of a test—SNR and sample size. Determining realistic ranges for both in the context of EEG FC estimation is challenging but critical. Second, due to source leakage, we must expect (tiny) spill-over effects from interacting to non-interacting region pairs, an effect termed “ghost interactions” (Palva et al., 2018). As a result, these ghost interactions will inevitably become statistically significant for any source pair at high enough SNRs and sample sizes—an effect that can also be seen in Figure 5c. For these reasons, we here assessed the effect sizes of FC metrics instead of than their statistical significance, and focused on evaluating the performance of different FC estimation pipelines relative to one another rather than on their absolute performance. However, future studies should go one step further by systematically assessing statistical maps derived from connectomes using our results as building blocks.

### Limitations

While this study investigates a large range of processing pipelines, phase-to-phase FC metrics, and data parameters, it is far from being exhaustive. Other works have shown that many other parameters like channel density (Song et al., 2015), the location of interacting sources (Anzolin et al., 2019), data length (Astolfi et al., 2007; Van Diessen et al., 2015; Liuzzi et al., 2017; Sommariva et al., 2019), referencing (Van Diessen et al., 2015; Chella et al., 2016; Huang et al., 2017), and co-registration (Liuzzi et al., 2017) can influence FC detection. Besides, we here used the same head model for generating the sensor data and estimating the inverse solution. However, we expect worse performance when the head model has to be estimated, and previous work has shown that the quality of head model estimation also influences FC detection (Mahjoory et al., 2017). Likewise, there exist many other inverse solutions, like MNE, wMNE, LORETA, sLORETA, and MSP, just to name a few. Further, there also exist other types of dimensionality reduction techniques. For example, some works selected the source with the highest power within a region or the source that showed the highest correlation to the time series of other sources in the ROI to be representative for all time series of the ROI (Hillebrand et al., 2012; Ghumare et al., 2018). Others have presented a procedure of optimizing a weighting scheme before averaging all time series within a ROI (Palva et al., 2010, 2011).

We also did not investigate the effect of the number of epochs and the epoch length in this study. It has been shown that the number of epochs can introduce a bias for certain connectivity metrics (Vinck et al., 2010). This is the case for connectivity metrics that yield positive values only, like (absolute) coherence, the absolute value of the imaginary part of coherency, MIM, or MIC. For these metrics, for a fixed epoch length, a lower number of epochs will systematically lead to higher values of estimated connectivity, even under the null hypothesis of no interaction. This is due to the higher variance of the estimates for lower samples sizes, which turns into a positive bias when the absolute value is taken. Further, Fraschini et al. (2016) argued that also the epoch length may have an influence on FC estimation, where shorter epochs were found to introduce a positive bias on FC when the number of epochs was held constant. As a result, we recommend to use fixed numbers of epochs throughout a single experiment. This is of particular importance when the goal is to compare different groups or experimental conditions. As the set of coupling mechanism and corresponding FC metrics that have been proposed is huge, we deliberately constrained our analysis here to phase-phase coupling using a selection of metrics that have previously been shown to be robust to mixing artifacts (Nolte et al., 2004; Haufe et al., 2013; Ewald et al., 2012). In contrast, non-robust metrics have been shown to be prone to the spurious discovery of interactions (Nolte et al., 2004; Haufe et al., 2013; Bastos and Schoffelen, 2016; Brunner et al., 2016; Van de Steen et al., 2019). This was confirmed here again for absolute coherence and GC. For a detailed overview of the taxonomy of FC metrics we refer to the works of Bastos and Schoffelen (2016); Schoffelen and Gross (2019); Marzetti et al. (2019). Our results are obtained for intra-frequency phase–phase coupling, and make no claims about non-linear interaction metrics quantifying phase–amplitude or amplitude–amplitude coupling within or across frequencies (De Pasquale et al., 2010; Hipp et al., 2012; Colclough et al., 2015). Nevertheless, we expect that robust-to-volume conduction measures for these FC types would be required to obtain optimal performance.

A further limitation of simulation studies in general is that assumptions need to be made that are hard, if not impossible, to confirm. Here, our goal was to generate pseudo-EEG data comprising realistic effects of volume conduction using a physical model of a human head. In terms of the generated time series, we focused on alpha-band oscillations as carriers of the modeled interactions. By adding pink brain noise, uniformly distributed across the entire brain, as well as white sensor noise, we obtained simulated sensor-space EEG data that resemble real data in crucial aspects such as spectral peaks and the general 1/f shape of the power spectrum. On the other hand, numerous additional assumptions were made regarding the linear dynamics of the interacting sources, the conception of the interaction as a pure and fixed time delay, the focus on an interaction in the alpha band, the number of interactions, the signal-to-noise ratio, and the stationarity of all signal and noise sources. Several of these experimental variables were systematically varied to provide a comprehensive picture of the performance of each pipeline in a wide range of scenarios. The ranking of the pipelines’ performances was robust in all tested scenarios. However, a remaining question is how realistic the individual studied parameter choices are. Our simulated environment resembles a setting of task-related (ongoing) activity with few dominant active and interacting sources, as opposed to a resting-state setting with numerous equally active and interacting sources. Hincapiée et al. (2017) showed that connectivity estimation pipelines including beamformers perform well for point-like sources, whereas for extended cortical patches, MNE source estimation was found to be more accurate. In this study, we simulated point-like sources, which could lead to an overestimation of beamformer performance. Considering that FC analyses are predominantly performed on ongoing (including resting-state) activity, the assumption of having only a few interacting source pairs standing out against non-interacting background sources may be challenged. However, this assumption was made here for the practical purpose of enabling a comparison between approaches. Considering that FC analyses are predominantly performed on ongoing (e.g., resting-state) activity rather than averaged data, the assumptions of only few interacting source pairs standing out against non-interacting background sources with relatively high SNR can certainly be questioned. However, these assumptions were made here for the practical purpose of enabling a comparison between approaches rather than with the ambition of claiming real-world validity.

Future simulation studies should nevertheless strive to further increase the realism of the generated pseudo-EEG signals. In this regard, Anzolin et al. (2021) presented a toolbox that mimics typical EEG artifacts like eye blinks. We restricted ourselves here to using artificial time series designed to exhibit the specific properties assessed by the studied FC metrics; that is, time-delayed linear dynamics. In contrast, biologically inspired models such as the models implemented within the virtual brain toolbox (Sanz Leon et al., 2013) provide a richer portfolio of non-linear dynamics and thus are alternative ground truth models specifically when the goal is to validate non-linear FC metrics. The COALIA model (Bensaid et al., 2019), for example, has been used to mimick network activity in epilepsy for the purpose of validating FC estimates (Allouch et al., 2022). Further studies used the same model family to study the effect of parameters such as electrode density on FC estimates (Tabbal et al., 2022; Allouch et al., 2023). Similarly, Jirsa and Müller (2013) have used TVB to evaluate metrics of cross-frequency coupling. Overall, these studies provide complementary evidence that is largely aligned with our results, for example with respect to the superiority of robust connectivity metrics. The plausibility of several assumptions made by neural mass models has also recently been questioned (Pathak et al., 2022). Nevertheless, such models hold great promise as validation tools in the future.

Note in this respect that it was not our intention to propose a realistic model of EEG data or even the whole brain but simply to generate data that would allow us to test how well ROI-level FC can be reconstructed in the presence of volume conduction/source leakage. The types of FC we are interested here (directed and undirected linear FC) have been widely studied and popular metrics to infer these types of FC are known to be heavily affected by volume conduction (Nolte et al., 2004; Haufe et al., 2013). Hence, it was our intention to identify metrics and pipelines that have a high chance of reconstructing FC on the ROI level when signals are mapped to the EEG and back by realistic forward and inverse models. We deliberately do not address the question whether networks estimated using FC metrics provide a correct depiction of actual brain networks.

As a further limitation, our simulations are to some extent restricted to EEG data. However, it can be expected that, qualitatively, the results of this paper could be transferred to MEG data. MEG analyses also suffer from the source leakage problem (Pizzella et al., 2014; Colclough et al., 2016) and benefit from disentangling signal sources with source reconstruction (Marzetti et al., 2019; Schoffelen and Gross, 2019). Moreover, the same FC metrics are typically used in EEG and MEG analyses (Schoffelen and Gross, 2009, 2019). Nevertheless, differences exist, which would be worth studying. In contrast to EEG, which records secondary neuronal return currents, MEG records the magnetic field that is induced by electrical activity and arises in a circular field around an electric current (Hämäläinen et al., 1993). Therefore, MEG cannot record radial neuronal currents (Huang et al., 2007). This must be taken into account when estimating the inverse solution from the leadfield, i.e. it is advised to reduce the rank of the forward model from three to two by applying an SVD at each source location (Westner et al., 2021).

We here provide a simulation framework that is openly accessible by the community. Individual pipeline steps, but also simulated data can easily be replaced by other variants, following a plug-and-play principle. A such, we encourage readers to test aspects of the pipelines, other data, and other FC metrics not considered here.

## Conclusion

This work compared an extensive set of data analysis pipelines for the purpose of extracting directed and undirected functional connectivity between predefined brain regions from simulated EEG data. While several individual steps of such pipelines have been benchmarked in previous studies, we focused specifically on the problem of aggregating source-reconstructed data into region-level time courses and, ultimately, region-to-region connectivity matrices. Thereby, we close a gap in the current literature evaluating FC estimation approaches. We show that using non-robust FC metrics greatly reduces the ability to correctly detect ground-truth FC. Further, in our simulated pseudo-EEG data, the use of the eLORETA inverse solution also leads to worse FC detection performance than beamformers. Moreover, the use of inverse solutions that are frequency-specific, such as DICS, may hamper the correct identification of the directionality of interactions. Finally, unequal dimensionalities of signals at different ROIs may bias certain connectivity measures, such as MIC and MIM, degrading their ability to identify true interactions from a noise floor. Thus, dimensionality reduction techniques should be applied such that the number of retained signal components is the same for all regions. We expect that avoiding these pitfalls may enhance the correct interpretation and comparability of results of future connectivity investigations. FC pipelines that show promising results with our simulated pseudo-EEG data consist of beamformer or champagne source reconstruction, aggregation of time series within ROIs using a fixed number of strongest PCs, and using a robust FC metric like MIM or TRGC. To which scenarios these results can be generalized remains to be shown in further studies. In practice, low SNR, high numbers of interactions, and small interaction delays may, however, reduce the performance even of the best performing pipelines.

## Acknowledgements

This project was supported by the European Research Council (ERC) under the European Union’s Horizon 2020 research and innovation programme (Grant agreement No. 758985) and through Deutsche Forschungsgemeinschaft (DFG, German Research Foundation) – Project-ID 424778381 – TRR 295. We thank Tien Dung Nguyen for contributing to the development of the ROIconnect plugin. The computations for this work were partly run on the open Neuroscience Gateway cluster (Sivagnanam et al., 2013).

## Data and code availability

The code for the simulation can be found here: https://github.com/fpellegrini/FCsim. The code for the ROIconnect plugin can be found here: https://github.com/sccn/roiconnect. And the code for the minimal real data example here: https://github.com/fpellegrini/MotorImag. Data of the real data example are available upon request.

## Appendix A.

### ROIconnect toolbox

ROIconnect is a freely available open-source plugin to the popular MATLAB-based open-source toolbox EEGLAB for EEG data analysis. It adds the functionality of calculating region-wise power and inter-regional FC on the source level. Moreover, it provides functions to visualize power and FC. All functions can be accessed by the EEGLAB GUI or the command line. ROIconnect uses core EEGLAB functions for importing and preprocessing EEG data, and calculating the leadfield and source model: we refer users to other EEGLAB functions to preprocess data before applying ROIconnect functions. The ROIconnect plugin can be downloaded through github ^4^ or installed via the EEGLAB GUI extension manager.

#### Key features

The features of ROIconnect are implemented in three main functions:

`Pop_roi_activity`, `pop_roi_onnect`, and `pop_roi_connectplot`. pop roi activity takes an EEG struct containing EEG sensor activity, a pointer to a headmodel and a source model, the atlas name, and the number of PCs for dimensionality reduction as input. It then calculates a source projection filter (default: LCMV) and applies it to the sensor data. Power is then calculated with the Welch method for every frequency on the voxel time series and then summed across voxels within regions. The result is saved in `EEG.roi.source_roi_power`. To estimate region-wise FC, the `pop_ roi_activity` function reduces the dimensionality of the time series of every region by employing a PCA and selecting the strongest PCs (as defined in the input) for every region. The resulting time series are then stored in `EEG.roi.source_roi_data`.

`Pop_roi_connec` calculates FC between regions. It builds on the output of `pop_roi_activity`. That is, it takes the EEG struct as input, as well as the name of the FC metrics that should be calculated. The function calculates all FC metrics in a frequency-resolved way. That is, the output contains FC scores for every region–region–frequency combination. To avoid biases due to different data lengths, `pop_roi_connect` estimates FC for time windows (‘snippets’) of 60 sec length (default), which subsequently can be averaged (default) or used as input for later statistical analyses. The snippet length can be flexibly adjusted by the user. The output of this function is stored under the name of the respective FC metric under `EEG.roi`.

The `pop_roi_connectplot` function enables visualizing power and FC in the following modes:

- Power as region-wise bar plot.
- Power as source-level cortical surface topography.
- FC as region-by-region matrix.
- Net FC, that is, the mean FC from all regions to all regions, as cortical surface topography.
- Seed FC, that is, the FC of a seed region to all other regions, as cortical surface topography.

For plotting, a specific frequency or frequency band can be chosen by the user. For matrix representations, it is also possible to just plot one of the hemispheres or only regions belonging to specific brain lobes.

## Footnotes

1 https://github.com/arnodelorme/roiconnect

2 https://github.com/fpellegrini/FCsim

3 https://github.com/fpellegrini/MotorImag

4 https://github.com/arnodelorme/roiconnect

