## Supplementary_Material for "Identifying good practices for detecting inter-regional linear functional connectivity from EEG"

*Preprint submitted to NeuroImage* *March 13, 2023*

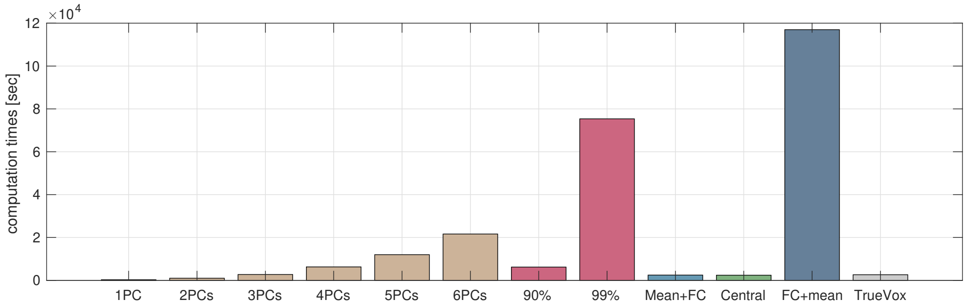


**Figure S1:** Comparison of computation times of all pipelines for Experiment 1B (single core, 16 GB allocated memory). Displayed is the summation of the aggregation within regions step and estimation of all FC metrics.

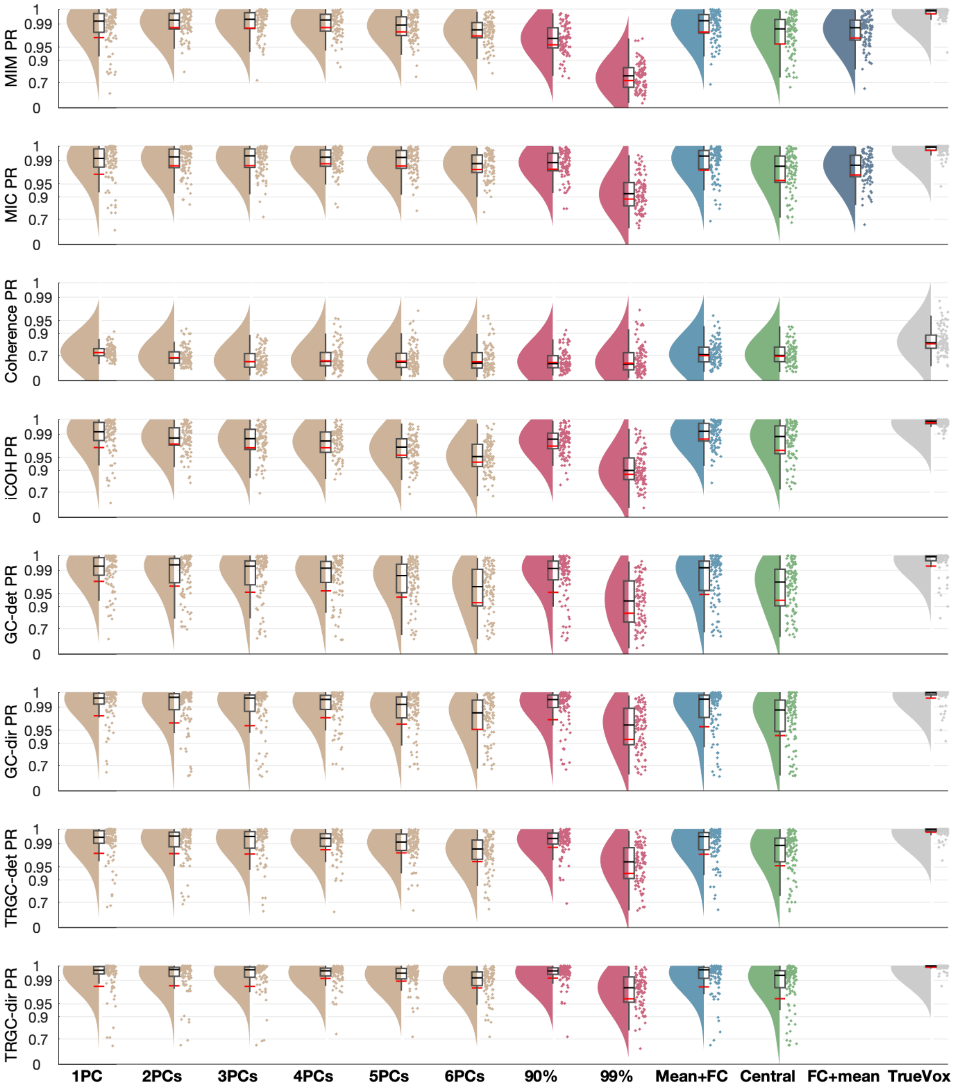


**Figure S2:** Full comparison of different pipelines and FC metrics (Experiment 1A and B). Red and black lines indicate the mean and median percentile rank (PR), respectively. The boxcar marks the 2.5th and 97.5th percentile.

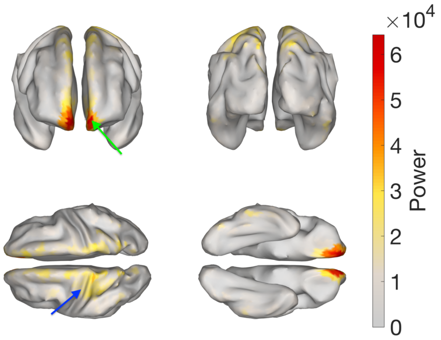


**(a)** Power map (eLORETA).

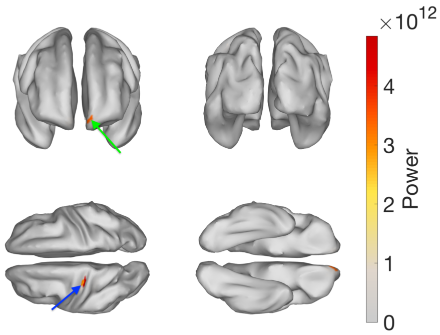


**(b)** Power map (LCMV).

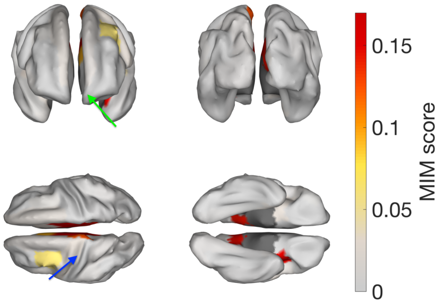


**(c)** MIM scores of seed region to all other regions (eLORETA).

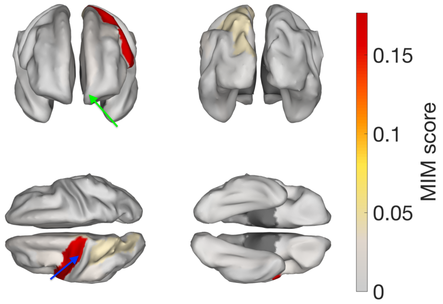


**(d)** MIM scores of seed region to all other regions (LCMV).

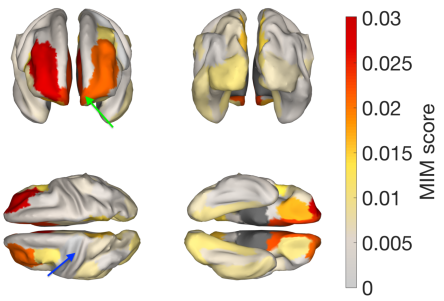


**(e)** MIM scores of target region to all other regions (eLORETA).

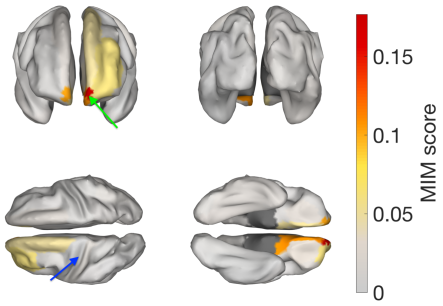


**(f)** MIM scores of target region to all other regions (LCMV).

**Figure S3:** Comparison of source localization with eLORETA vs. LCMV. Activity and ground-truth interaction between a seed and a target voxel was simulated, and the FIXPC1 pipeline was applied. We show the resulting power maps (a + b), and seed and target MIM scores (c to f). The green arrow points to the seed voxel, the blue arrow to the target voxel.

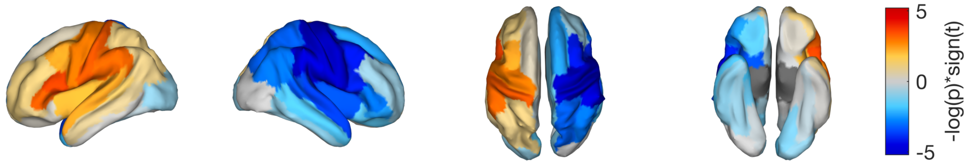


**Figure S4:** Comparison of power in left vs. right hand motor imagery tasks. Depicted are negative log10-transformed p-values, multiplied with the sign of the t-statistic. Positive values indicate higher power in the left hand motor imagery trials.
